# Working Memory as Programmable Fast Weight Computation

**DOI:** 10.64898/2026.06.02.729709

**Authors:** Longsheng Jiang, Yanlin Zhu, Jia Liu

## Abstract

Working memory (WM) stores information after sensory input disappears and later retrieves it in a task-relevant format, but the mechanism unifying storage and retrieval remains unclear. Here we combine neural geometry analyses of macaque dorsolateral prefrontal cortex activity during a visuospatial delayed-match-to-sample task with computational modeling to test whether WM can be implemented as recurrent fast-weight computation. We found that the relational geometry of remembered locations was strongly expressed during sample presentation, degraded during the early delay, and reemerged before requirement in a partially distinct mnemonic subspace. A recurrent fast-weight programmer model, which implements a form of dynamic fast-weight memory closely related to linear Transformer computation, reproduced these latent-to-mnemonic dynamics. Direct inspection and perturbation of the model revealed that neural activity writes stimulus information into rapidly modifiable synaptic states, synaptic dynamics organize this latent memory over time, and recurrent readout queries the evolving state to generate task-relevant activity. These findings provide a unified account of WM storage and retrieval and suggest that biological WM and Transformer family architectures share an algorithmic principle of programmable temporary memory.

## Introduction

Working memory (WM) converts a fleeting sensory event into an internal state that can be held, reorganized, and later used to guide behavior. This process contains two inseparable operations. Information must be stored after the external stimulus disappears, and it must later be retrieved in a format appropriate for the downstream computation. Existing theories have often emphasized one operation more than the other. Persistent-activity theories explain how information can remain expressed in neural activity during a delay (Funahashi et al., 1989; Fuster & Alexander, 1971), whereas synaptic theories explain how information can be stored when stimulus-specific neural firing is weak or absent (Mongillo et al., 2008; Stokes, 2015; Lundqvist et al., 2018). However, how the retrieval occurs remains insufficiently explained.

The retrieval problem is particularly important because WM is expressed dynamically rather than statically. In many delayed-response tasks, neural and behavioral signals gradually increase before the remembered information is needed, demonstrated as ramping firing-rate activity (Rainer & Miller, 2002; Warden & Miller, 2010; Tian et al., 2024), oscillatory bursts (Lundqvist et al., 2016), evolving population codes (Barak et al., 2010a; Inagaki et al., 2019; Spaak et al., 2017), and memory-related micro-gaze patterns (Linde-Domingo & Spitzer, 2023). These observations raise a mechanistic question: How does information stored in latent synaptic states become reinstated as anticipatory neural activity? One possibility is that storage and retrieval result from a coherent trajectory through synaptic state space. Transient neural activity writes information into rapidly modifiable synaptic states; these states preserve the memory while stimulus-specific neural activity becomes weak or absent; and ongoing synaptic dynamics then make the stored content progressively readable in a task-relevant neural format.

Testing this hypothesis empirically is challenging because synaptic states are not directly accessible in standard neurophysiological recordings. Instead, computational models can expose and perturb synaptic-like variables, and previous modeling work has shown that short-term synaptic plasticity can support activity-silent maintenance (Mongillo et al., 2008; Rolls et al., 2013; Mi et al., 2017; Pals et al., 2020; Lansner et al., 2023). However, many such models focus mainly on storage and often retrieve the memory in the same representational format in which it was encoded. Real WM behavior often requires more than reinstating a past sensory pattern; indeed, a remembered stimulus may need to be transformed into a decision variable (Panichello & Buschman, 2021), an action plan (Nasrawi et al., 2023), or a mnemonic code protected from interference with new sensory input (Libby & Buschman, 2021). This functional demand requires a model that can maintain latent content, reorganize it across the delay, and retrieve it into a new neural subspace.

Fast-weight programming provides a natural computational framework for this purpose. In a fast-weight programmer (FWP), one network rapidly modifies the weights of another network, creating a temporary memory state that can later be queried (Schmidhuber, 1992). This architecture is intrinsically similar to synaptic theories of WM, as neural activity can program short-lived changes in synaptic efficacy, and subsequent activity can read from these modified synaptic states. Thus, FWP provides an algorithmic model of programmable temporal memory. Interestingly, FWP also links WM to the Transformer family, a central architecture family in modern AI. Fast weights and dynamic connections have long been explored in AI as mechanisms for rapidly modifying memory states (Feldman, 1982; Hinton & Plaut, 1987). More recently, linear Transformers, which are mathematically equivalent to FWPs, have been shown to instantiate this principle: key-value outer products construct a dynamic matrix-valued memory, and queries read from this memory during sequence processing (Schlag et al., 2021; Irie & Gershman, 2025). From WM perspective, this matrix is a temporary memory buffer. This equivalence suggests that biological WM and the Transformer family may share an approximate algorithmic principle based on fast programmable memory states.

Here we tested whether this fast-weight computation underlies the WM process from storage to retrieval. In a combined empirical and modeling strategy, neural recordings defined the population-level phenomenon that a model must explain, while modeling tested whether a specific synaptic mechanism was necessary and sufficient to generate that phenomenon. Specifically, we first analyzed macaque dorsolateral prefrontal cortex (dlPFC) neural activity recorded during a visuospatial delayed-match-to-sample (DMS) task (Meyer et al., 2011; Qi et al., 2011; Tang et al., 2022), and examined whether the relational geometry among stimulus locations was preserved, degraded, or recovered across the delay. We then developed a recurrent FWP model trained on a minimal representation-transfer task designed to capture the sensory-to-mnemonic transformation. This model allowed us to ask whether rapidly updated synaptic-like states could account for the mechanism that store latent information and retrieve it into an expressed mnemonic code.

## Results

### Memory Demand Reveals Late Recovery of a Degraded Stimulus Code

We first reanalyzed dlPFC activity recorded while monkeys performed a visuospatial DMS task (Fig. 1a; see Methods) (Meyer et al., 2011; Qi et al., 2011). On each trial, the monkey fixated centrally while a white square appeared at one of nine locations, in which the central location was the reference and the eight peripheral locations were the spatial stimuli. The sample was followed by a first delay, a probe stimulus, a second delay, and a response period (Fig. 1b). The task included passive and active conditions with matched sensory events but different memory demands. In the passive condition, monkeys maintained fixation and received reward without making an explicit judgment. In the active condition, monkeys judged whether the probe appeared at the sample location or at the diametrically opposite location, and reported the choice with a saccade. Only correct judgments were rewarded. Thus, the active condition required retention of the sample location across the first delay, whereas the passive condition provided a sensory and fixation control without an explicit WM requirement. To isolate activity related to sample encoding and maintenance, we restricted analyses to fixation, sample presentation, and the first delay, before probe onset and decision-related processing.

**Figure 1.**
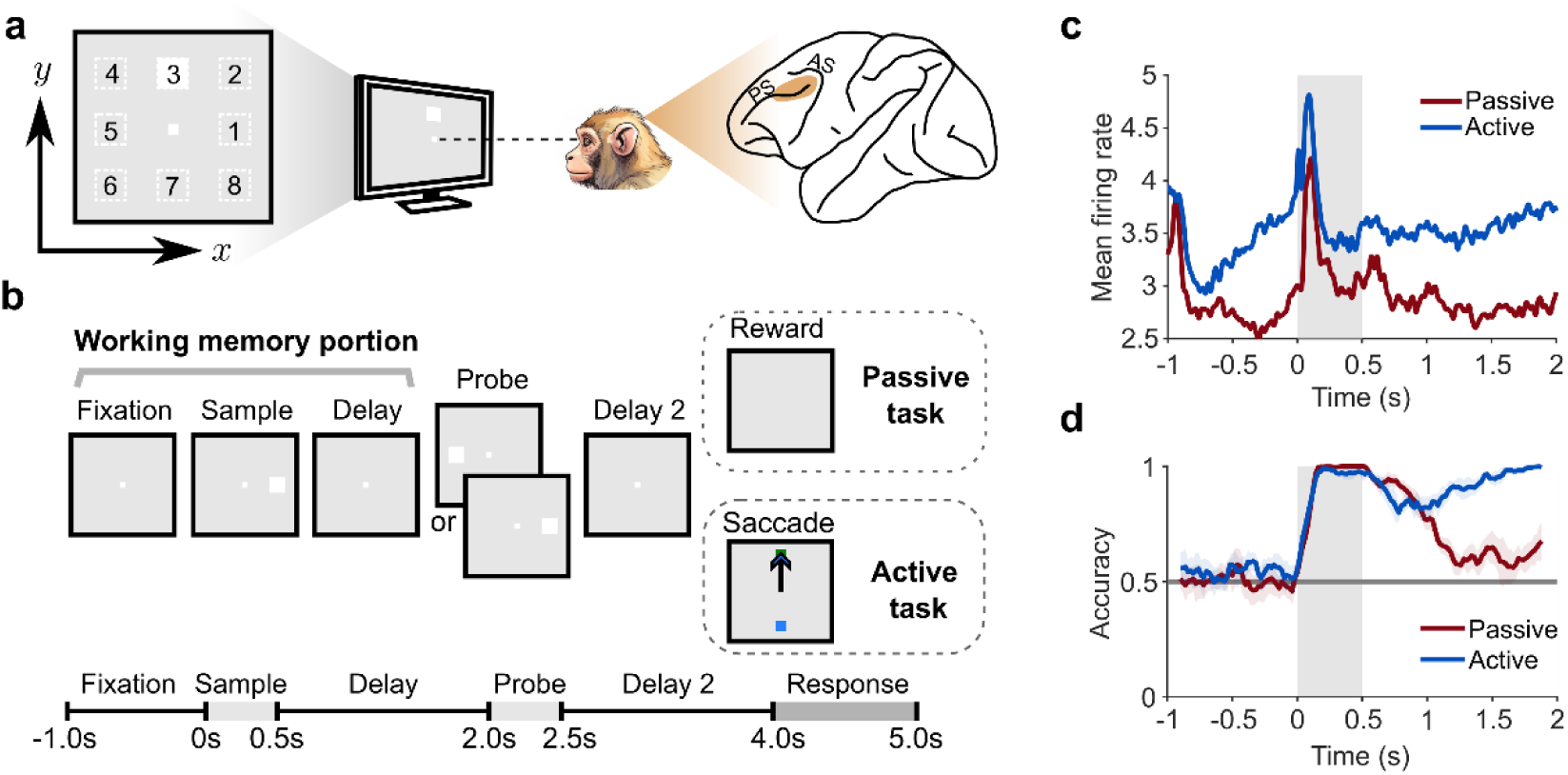
Task design and initial dlPFC population dynamics. **(a)** Schematic of the visuospatial DMS task. Left: The monkey fixated centrally while a peripheral sample stimulus (white square) appeared at one of eight possible locations. Right: Lateral view of the monkey brain showing the dlPFC recording site above the principal sulcus (PS) and near the arcuate sulcus (AS). **(b)** Trial timeline. The fixation, sample presentation, and first delay constituted the WM portion analyzed here. In the passive condition, monkeys maintained fixation and received reward without an explicit judgment. In the active condition, the monkey reported whether the probe matched the sample by making a saccade during the response period. **(c)** Population-averaged firing rates of dlPFC neurons in the example animal (Monkey ELV), aligned to sample onset for the active (blue) and passive (red) conditions. Gray region indicates sample presentation. **(d)** Time-resolved decoding accuracy for discriminating diametrically opposite sample locations. The horizontal gray line indicates chance performance. Colored shading indicates standard deviation (SD) across decoding analyses.

Population-averaged firing rates provided an initial view of delay-period dynamics (Fig. 1c). In the active condition, dlPFC activity showed a transient stimulus-evoked response, followed by a reduction during the early (0.5–1.0 s) and middle delay (1.0–1.5 s), and then a modest increase toward the end of the delay (1.5–2.0 s). In the passive condition, sample presentation also evoked activity, but firing returned to a lower baseline and showed no comparable late-delay increase. Thus, mean firing rate revealed only a weak ramping signal that depended on memory demand, consistent with previous studies (Romo et al., 1999; Rainer & Miller, 2002; Barak et al., 2010b; Warden & Miller, 2010). This univariate measure alone could not determine whether information about the sample location was lost, maintained silently, or reformatted during the delay.

We therefore asked whether stimulus identity remained detectable in the multivariate population pattern. Linear classifiers were used to quantify the time-resolved separability of diametrically opposite sample locations, such as locations 1 versus 5 (Fig. 1d; see Methods). During sample presentation, decoding rapidly increased to near-ceiling accuracy in both active and passive conditions, indicating strong sensory encoding of sample location. After sample offset, decoding in the passive condition declined gradually toward chance, consistent with the absence of an explicit requirement to retain the sample. In contrast, decoding in the active condition initially weakened after sample offset but subsequently recovered during the delay, reaching near-ceiling accuracy by the end of the delay. This dissociation between conditions identifies a memory-demand-dependent recovery of the dlPFC population code (Stokes, 2015). Next, to look beyond unstructured stimulus identity, we tested whether the recovered code preserved low-dimensional geometric spatial relationships of the stimuli and whether it differed from the initial sensory code.

### Spatial Relationship Becomes Latent and Reemerges During Working Memory

Because the eight sample locations were arranged on a circle, the task provided a natural physical geometry against which dlPFC population geometry could be evaluated (Fig. 2a). Thus, we asked whether population activity preserved the geometrical structure of the sample locations. This geometric approach is well suited for characterizing structured WM representations (Xie et al., 2022; Pu et al., 2024; Ma et al., 2025), because it tests whether neural activity maintains the organization of the stimulus space.

**Figure 2.**
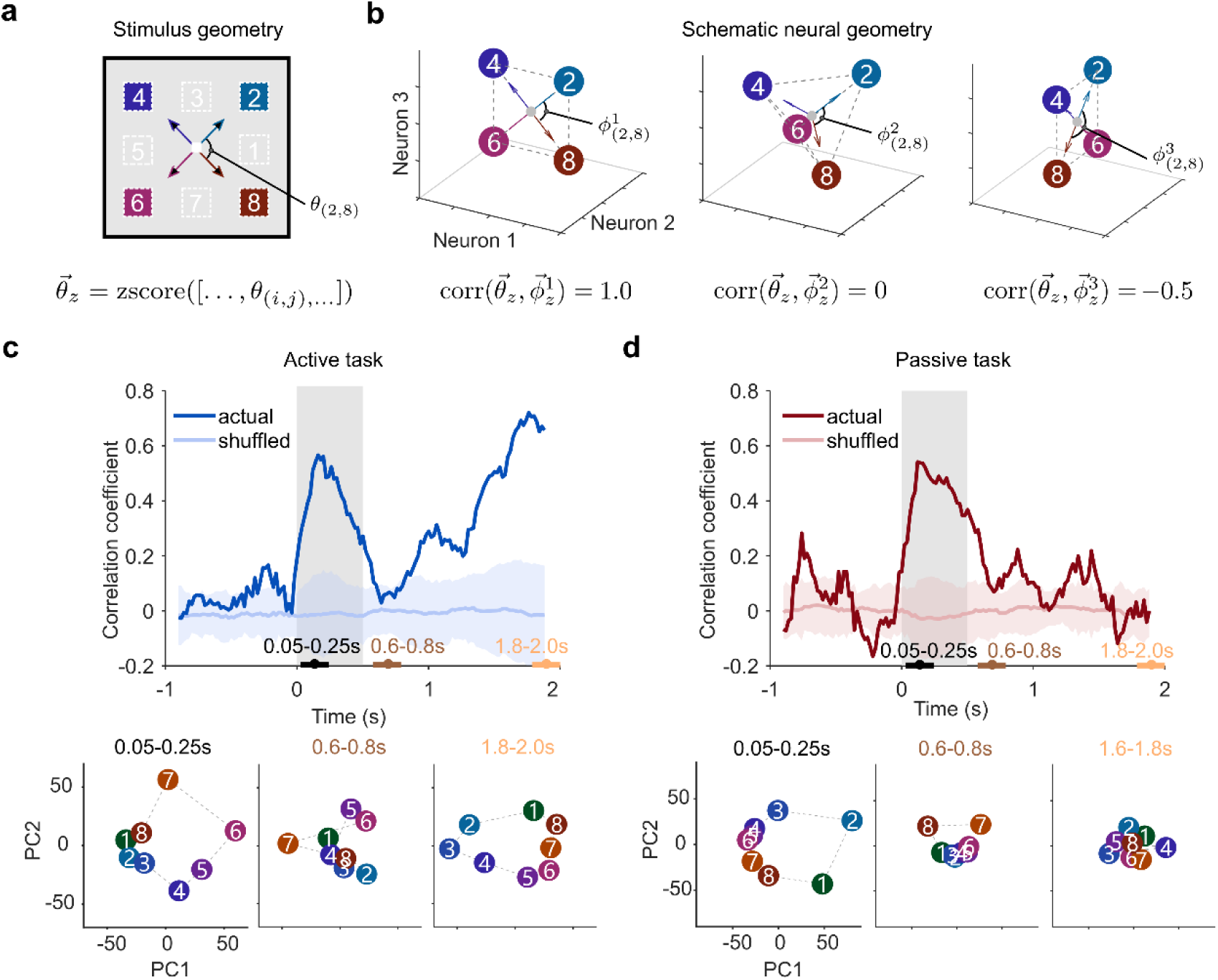
Neural Geometry Analysis of Spatial Relationship Coding. **(a)** Physical stimulus geometry. Four example locations are shown from the full set of eight circularly-arranged locations. Vectors from fixation to the stimulus locations define the physical stimulus geometry. Pairwise angles between vectors (𝜃_(𝑖,𝑗)_, 𝑖, 𝑗: two locations) define the stimulus aRDV 𝜃⃗ = [𝜃_(𝑖,𝑗)_], 𝑖 < 𝑗. 𝜃⃗_𝑧_ denotes z-scored 𝜃⃗. **(b)** Schematic examples of neural geometries. At each time point, the population response to each stimulus was represented as a vector from the state of central reference location to the corresponding stimulus-specific neural state. 𝜙⃗⃗^𝑘^ denotes the aRDV of 𝑘th exemplar neural geometry (𝑘 = 1, 2, 3), and 𝜙⃗⃗^𝑘^ represents z-scored 𝜙⃗⃗^𝑘^. Left: A neural geometry that preserves the stimulus geometry and yields a high positive stimulus-neural geometry correlation. Middle: A neural geometry unrelated to the stimulus geometry and yielding a near-zero correlation. Right: A neural geometry inversely related to the stimulus geometry and yielding a negative correlation. **(c)** Active condition. Top: Time-resolved stimulus-neural geometry correlation. Gray region denotes the sample presentation period. The light blue shading denotes the shuffled-label baseline, shown as mean ± SD across 100 repetitions. Bottom: PCA visualizations of neural states in the sample presentation, early-delay, and late-delay periods. **(d)** Passive condition. Same conventions as in (c).

At each time point, the population response to each sample location was represented as a vector in a high-dimensional neural state space, with one dimension per neuron. The relative neural vector for each stimulus location was defined as the displacement from the state of the central reference location to the corresponding stimulus-specific population state. We then computed the pairwise angles between all stimulus-specific neural vectors to obtain a neural angular representational dissimilarity vector, referred to as the neural aRDV (Fig. 2b; See Methods). In parallel, we computed an analogous stimulus aRDV from the pairwise angles between physical stimulus vectors in the two-dimensional display space. The correlation between the z-scored neural aRDV and the z-scored stimulus aRDV quantified the extent to which the neural population preserved the circular spatial geometry of the samples. High correlations indicated that angular relationships among locations in physical space were reflected in neural state space; and near-zero correlations indicated that this spatial geometry was not expressed in the population activity.

In the active condition, the stimulus-neural geometry correlation rose rapidly after sample onset, indicating that dlPFC activity formed a population geometry aligned with the physical geometry (0.05–0.25 s, mean = 0.49; Fig. 2c, top). This stimulus-aligned geometry was evident in the space spanned by the top two principal components from principal component analysis (PCA). The neural states formed an approximately circular arrangement that matched the layout of the stimulus locations (Fig. 2c, bottom left). This PCA projection was used only for visualization, whereas the geometry correlation was computed from the high-dimensional neural vectors.

After sample offset, the geometry correlation declined to the baseline during the early delay (0.6–0.8 s, mean = 0.06; permutation test, 𝑝 = 0.25). In the corresponding PCA projection, the neural states no longer formed an ordered circular structure and instead collapsed into an unstructured cloud (Fig. 2c, bottom middle). Because the preceding decoding analysis showed that some stimulus-discriminative information could remain detectable during this early delay, this collapse should not be interpreted as a complete loss of all stimulus-related information. Rather, it shows that the structured spatial relationship among locations was no longer accessible to the population-geometry measure, consistent with an operationally activity-silent or latent state.

During the late delay, the geometry correlation gradually increased and became significant before probe onset (1.8–2.0 s, mean = 0.68; permutation test, 𝑝 < 0.01). This correlation was comparable with that in sample presentation when no windowed smoothing was applied (Supplementary Fig. 3). The corresponding PCA projection revealed a reemergent circular arrangement of neural states, indicating that the spatial relationship among stimulus locations was reinstated in population activity (Fig. 2c, bottom right). Thus, under active WM demand, dlPFC activity recovered a structured representation of the spatial relationship defined by the sample locations.

This degradation-and recovery pattern was also observed when using distance-based RDV (Supplementary Fig. 4). It might weaken when using smaller neural population (Supplementary Fig. 5) and similar pattern was observed in the other two monkeys (Supplementary Fig. 6).

In contrast, this degradation-and-recovery dynamic was absent in the passive condition. During passive viewing, sample onset again produced a transient increase in the geometry correlation (0.05–0.25 s, mean = 0.48), indicating that the sensory-driven population response initially reflected the spatial layout of the samples (Fig. 2d, top). However, after sample offset, the correlation returned to the baseline and did not recover during the delay (0.6–0.8 s, mean = 0.12; 1.8–2.0 s, mean = -0.01). The corresponding PCA projections showed a stimulus-aligned circular geometry during sample presentation, but no clear spatial structure in either the early or late delay (Fig. 2d, bottom).

Together, these results show that WM demand selectively induced the degradation and subsequent recovery of spatial relational geometry in dlPFC population activity, providing a population-level signature of latent memory maintenance and active retrieval. This dynamic raises the question of whether the recovered geometry returned to the original sensory subspace or was reconstructed in a distinct mnemonic subspace. This question is important because a neural population that must maintain past information while processing new inputs may benefit from separating sensory and mnemonic codes, thereby reducing interference between incoming stimuli and remembered content (Libby & Buschman, 2021).

### Spatial Geometry Reemerges in a Partially Orthogonal Mnemonic Subspace

To examine whether dlPFC population activity separates sensory and mnemonic codes, we performed cross-temporal decoding analysis (Cavanagh et al., 2018; Spaak et al., 2017). For each pairwise combination of the eight peripheral sample locations, we trained a linear classifier at one time point and tested it at all other time points across a trial, yielding 28 pairwise decoders. Strong generalization, when information is decodable at both time points, indicates that stimulus information is represented along similar coding axes. Weak generalization, on the other hand, indicates that the representation has been transformed into a different neural subspace.

We first used the probe as a sensory benchmark. In match trials, the sample and probe appeared at the same location; as expected, a decoder trained during sample presentation generalized strongly to probe presentation (2.05–2.45 s) in both the passive (mean=0.95; Fig. 3a) and active conditions (mean = 0.85; Fig. 3b; see Methods). In nonmatch trials, in which the probe appeared at the diametrically opposite location, the corresponding decoder outputs were inverted (Supplementary Fig. 7). These results indicate that repeated sensory information occupied a largely shared sensory subspace.

**Figure 3:**
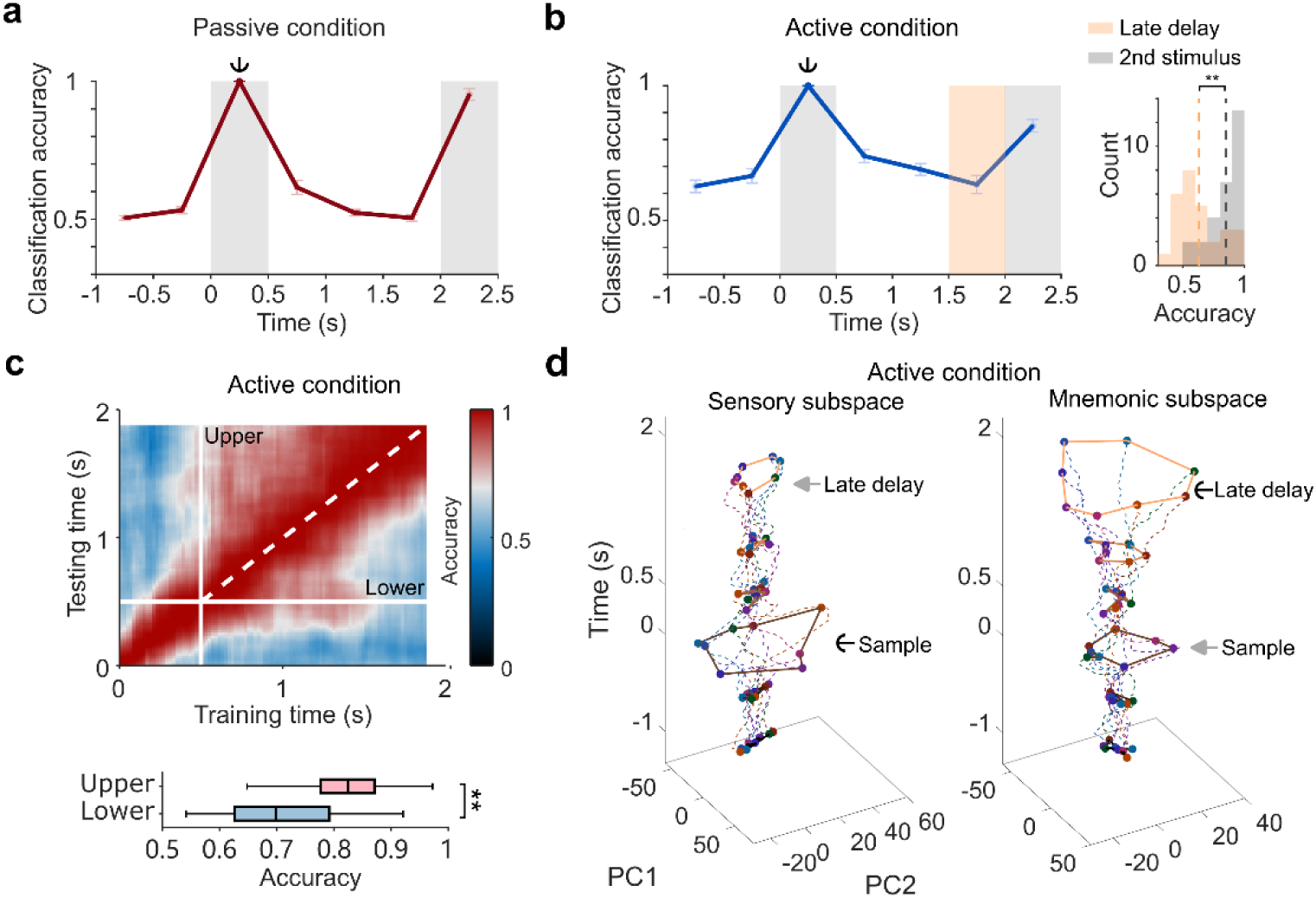
Sensory and mnemonic spatial codes occupy partially distinct neural subspaces. **(a)** Cross-temporal decoding in the passive condition. Linear classifiers were trained during sample presentation, indicated by the anchor, and tested across the trial in match trials. **(b)** Cross-temporal decoding in the active condition. Left: Same conventions as in (a). Orange shading denotes the late delay. Right: Distribution of decoding accuracies across the 28 decoders when tested during the late delay and during probe presentation. Dashed lines denote mean accuracies. **(c)** Cross-temporal decoding matrix in the active condition. Top: Each pixel in the matrix shows the mean accuracy of the 28 decoders trained at the time on the x-axis and tested at the time on the y-axis. White lines indicate sample offset, and the dashed line indicates the matrix diagonal. Bottom: Accuracy distributions in the upper and lower triangular regions. **(d)** Cross-temporal PCA projections in the active condition. Neural states were projected onto a sensory subspace defined from sample-period activity (Left) or a mnemonic subspace defined from late-delay activity (Right). Both subspaces preserved the circular organization of sample locations, but with different native coding coordinates. **: 𝑝 < 0.001.

The late-delay mnemonic code in the active condition showed a different pattern. Decoders trained during sample presentation generalized only moderately to the late delay (mean = 0.63; chance = 0.5), significantly less than their generalization to probe presentation (Wilcoxon signed-rank test, 𝑝 < 0.001; Fig. 3b). Specifically, approximately half of the 28 pairwise decoders showed late-delay accuracies clustered near chance (0.4–0.6; Fig 3b, right). Because the preceding analyses showed that both spatial identity and relational geometry reemerged during the late delay, this reduced cross-temporal generalization cannot be attributed simply to a loss of mnemonic information. Instead, it suggests that the remembered spatial structure was recovered in a coding space that differed from the initial sensory space.

To test this intuition, we further characterized the temporal evolution of the delay-period code using a full cross-temporal decoding matrix in the active condition (Fig. 3c; see Methods). Within the delay period, decoding accuracy was higher in the upper triangular region of the matrix, where decoders were trained at earlier time points and tested at later time points, than in the lower triangular region, where decoders were trained later and tested earlier (t-test, 𝑝 < 0.001). This asymmetry indicates that the delay-period code evolved along a directed trajectory: earlier delay states were more compatible with later delay states than later delay states were with earlier delay states. Thus, the delay-period representation did not occupy a static coding space; instead, it gradually transformed from a sensory subspace toward a late-delay mnemonic subspace.

To visualize this transformation, we performed cross-temporal PCA projection analysis. We defined a sensory subspace from sample-evoked activity and a mnemonic subspace from late-delay activity. Neural states from all time points were then projected onto the top two principal components of each subspace (see Methods). In the active condition, native projections revealed stimulus-aligned circular geometries in both subspaces. Sample-period activity formed an expanded circular structure in the sensory subspace (area = 3538, arbitrary unit, a.u.), and late-delay activity formed an expanded circular structure in the mnemonic subspace (area = 3439 a.u.; Fig. 3d). Cross projections between the two subspaces preserved the circular structure but reduced its scale. Late-delay activity projected into the sensory subspace formed a compressed geometry (area = 720 a.u.), and sample-period activity projected into the mnemonic subspace was similarly compressed (area = 814 a.u.). These results indicate that the sensory and mnemonic codes represented the same spatial structure in partially distinct neural subspaces.

Together, the cross-temporal decoding and projection analyses show that the remembered spatial geometry reemerged in a mnemonic subspace that was distinct from, but not orthogonal to, the sensory subspace. This finding raises two mechanistic questions. First, where was memory information stored during the early delay, when spatial geometry was not expressed in neural activity? Second, how was the latent representation gradually reconstructed in a distinct mnemonic subspace? Synaptic theories (Mongillo et al., 2008; Mi et al., 2017; Mongillo, 2025; Mongillo & Tsodyks, 2026) propose that maintenance and reconstruction may both depend on rapidly changing synaptic states. Therefore, we turned to computational modeling to test whether dynamic synaptic-like states could account for the storage and retrieval of the observed population geometry.

### A Recurrent Fast-Weight Model Reproduces Latent-to-Mnemonic Geometry Recovery

To examine whether the WM dynamics observed in dlPFC could arise from dynamic synaptic-weight updating, we built a recurrent fast-weight programmer model (FWP; Fig. 4a). Classical FWPs consist of a slow net that programs the rapidly changing weights of a fast net (Schmidhuber, 1992), and recent work has shown that linear Transformers can be interpreted as a feedforward special case of this broader fast-weight framework (Schlag et al., 2021). In our model, the fast net was a recurrent neural circuit whose hidden activity was treated as model population activity, while its dynamic recurrent weights were treated as synaptic-like state variables. The slow net received the same external input as the fast net and generated update signals for these dynamic recurrent weights. The update rule of the slow net contained two complementary components. A Hebbian-like associative term allowed stimulus-evoked activity patterns to be written into the fast weights (Schmidhuber, 1992), and a drift term allowed the fast weights to evolve along a directed delay-period trajectory (Seeholzer et al., 2019). Thus, the model implements short-term synaptic plasticity at the algorithmic level, without assuming a specific biophysical mechanism for synaptic dynamics (Lundqvist et al., 2011).

**Figure 4:**
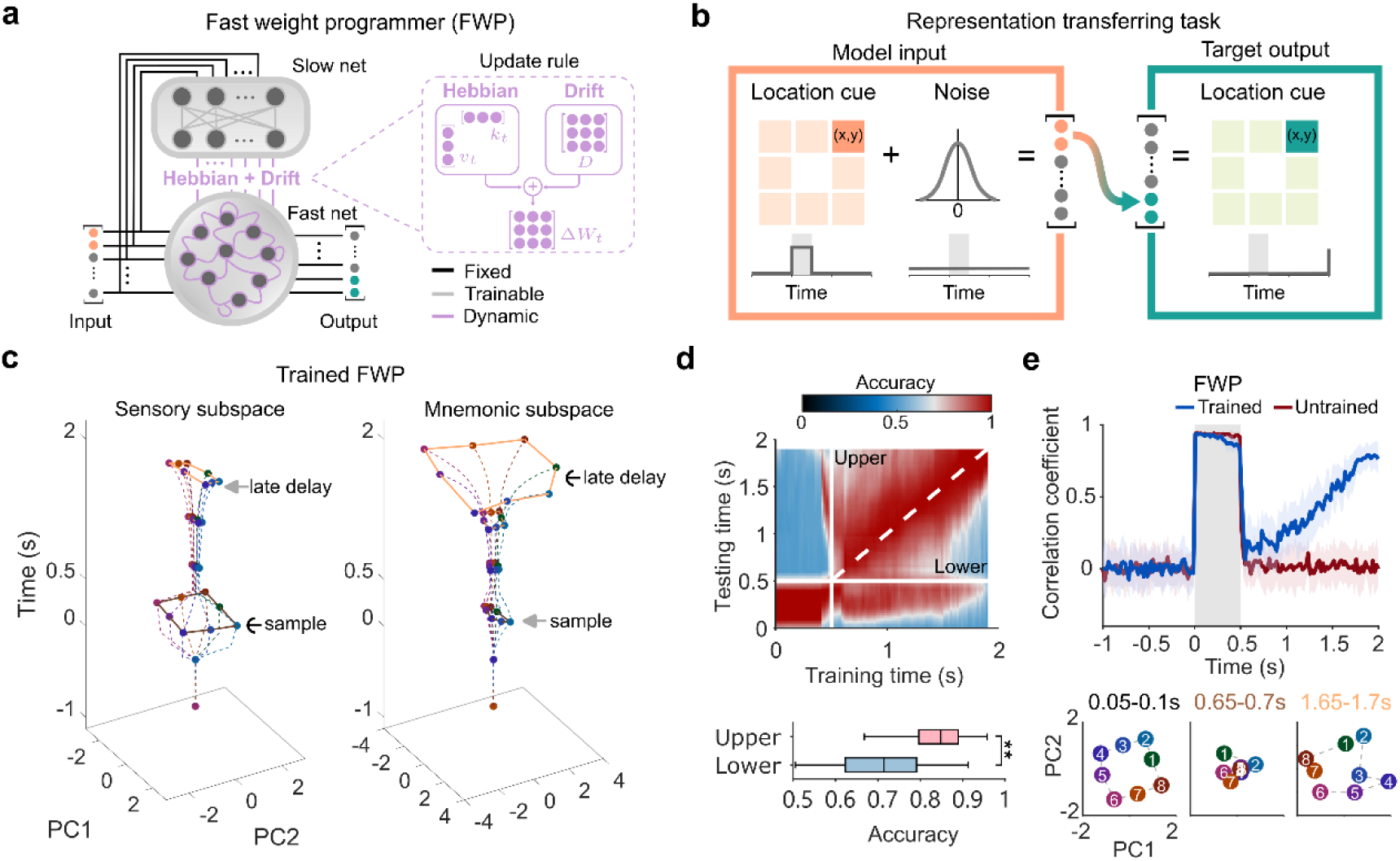
A recurrent FWP model reproduces latent-to-mnemonic geometry recovery. **(a)** Architecture of the recurrent FWP model. The model consists of a slow net and a recurrent fast net. The fast net generates population activity and output. Its recurrent weights are dynamically updated by an update signal Δ 𝑊_𝑡_ which combines a Hebbian component and a drift component. The Hebbian term is the outer product of a key vector 𝑘_𝑡_ and a value vector 𝑣_𝑡_, and the drift term is a constant matrix 𝐷, where 𝑘_𝑡_, 𝑣_𝑡_, and 𝐷 are from the slow net. Black, gray, and purple connections denote fixed, trainable, and dynamic weights, respectively. **(b)** Representation-transfer task. A transient two-dimensional stimulus cue is embedded in the first input dimensions against trial-long Gaussian noise. The target output is the same cue expressed in the last output dimensions and is evaluated only at the end of the delay. **(c)** Cross-temporal projections of trained fast-net activity into the sensory subspace (left) and the mnemonic subspace (right). Stimulus-period activity is expanded in the sensory subspace, whereas late-delay activity is expanded in the mnemonic subspace. **(d)** Cross-temporal decoding matrix for fast-net activity. White lines indicate stimulus offset, and the dashed line indicates the matrix diagonal. Upper and lower denote the upper-left and lower-right triangular regions, respectively. The bottom panel shows decoding accuracies in the upper and lower triangular regions of the delay-period matrix. **(e)** Time-resolved stimulus-neural geometry correlation for trained and untrained models. Gray region denotes sample presentation, and colored shading denotes the SD across 20 repetitions. The bottom panels show neural geometries sampled in sample presentation, early delay, and late delay in one trial. The layouts in sample presentation and late delay were aligned with the stimulus layout. **: 𝑝 < 0.001.

We trained the model on a minimal representation-transfer task designed to capture the empirical transition from a sensory code to a mnemonic code (Fig. 4b; see Methods). On each trial, the input consisted of a transient two-dimensional stimulus cue embedded in ongoing Gaussian noise. The cue was presented only during the sample period and was injected into the first two dimensions of the input space, whereas noise was present throughout the trial and spanned all input dimensions. The target output was the same two-dimensional cue, but expressed in the last two dimensions of the output space. Importantly, the target was imposed only at the end of the delay. Thus, the model was trained to solve a memory-transfer problem, not to reproduce the empirical degradation-and-recovery dynamic. Internal dynamics were assessed only after training, using the same geometry and cross-temporal analyses applied to the dlPFC data.

After training, the model successfully performed the representation-transfer task (Supplementary Fig. 8; see Methods). As expected, cross-temporal projection analysis showed that stimulus-period activity formed an expanded stimulus-aligned structure when projected into the sensory subspace, whereas late-delay activity was compressed in that subspace (Fig. 4c, left). Conversely, late-delay activity formed an expanded stimulus-aligned structure when projected into the mnemonic subspace, whereas stimulus-period activity was compressed in that subspace (Fig. 4c, right). These projections indicate that the model preserved the relational structure of the stimulus space while transforming the neural coordinates in accordance with task demanded.

Cross-temporal decoding of fast-net activity further revealed that delay-period decoding was significantly stronger in the upper-triangular region of the cross-temporal matrix than in the lower triangular region (t-test, 𝑝 < 0.001; Fig. 4d), indicating a forward-evolving delay-period code. Thus, as in the dlPFC data, the model representation gradually moved away from the sensory state and approached a late mnemonic state. This transformed activity reproduced the degradation-and-recovery of spatial geometry. During sample presentation, the stimulus-neural geometry correlation rose rapidly, indicating that fast-net population activity initially formed a geometry aligned with the stimulus layout (0.05–0.25 s; mean = 0.93; Fig. 4e). After sample offset, this correlation declined close to baseline during the early delay, indicating that the stimulus layout was only weakly expressed in fast-net activity (0.6–0.8 s; mean = 0.17). During the late delay, however, the correlation gradually increased, indicating recovery of the stimulus-aligned geometry before the time at which the output was required (1.8–2.0 s; mean = 0.78). This latent-to-active recovery was absent in an untrained model with the same architecture and randomly initialized trainable parameters (sample presentation, 0.05–0.25 s, mean = 0.96; early delay, 0.6–0.8 s, mean = 0.04; late delay, 1.8–2.0 s, mean = 0.01; Fig. 4e). Thus, the late recovery of stimulus geometry required task-optimized fast-weight updating and was not a trivial consequence of the network architecture.

Together, these results show that the recurrent FWP model trained to solve a representation-transfer task spontaneously reproduced the empirical degradation-and-recovery dynamics observed in dlPFC. Next, we performed a series of ablation analyses to determine which model components were required to reproduce this neural signature.

### Hebbian Writing, Synaptic Drift, and Recurrence Are Jointly Required

We first examined whether fixed recurrent dynamics, in the absence of fast-weight control from the slow net, could generate the same degradation-recovery dynamics. Specifically, we trained a conventional recurrent neural network (RNN) with the same hidden-sate architecture as the fast net, except that its recurrent weights were fixed after training and received no dynamic updates from a slow net (Fig. 5a, left). The trained RNN successfully performed the representation-transfer task (Supplementary Fig. 9; see Methods). However, its internal dynamics differed qualitatively from both the dlPFC data and the recurrent FWP results. The RNN expressed a stimulus-aligned geometry continuously across the delay (0.5–2.0 s; mean = 0.87; Fig. 5a, right). Thus, the RNN solved the task by maintaining an activity-expressed population code, rather than by storing the stimulus in a latent state and reconstructing it later. Supplementary cross-temporal analyses further revealed that the RNN transformed its coding subspace through rotational dynamics (Supplementary Fig. 10). A more biologically constrained RNN incorporating excitation-inhibition balance and synaptic-efficacy-gated recurrent connections (Masse et al., 2019) also failed to reproduce the empirical degradation-recovery dynamics (Supplementary Fig. 11). These results indicate that, within the model classes tested, recurrent neural dynamics alone were insufficient to reproduce the latent-to-mnemonic recovery observed in dlPFC.

**Figure 5:**
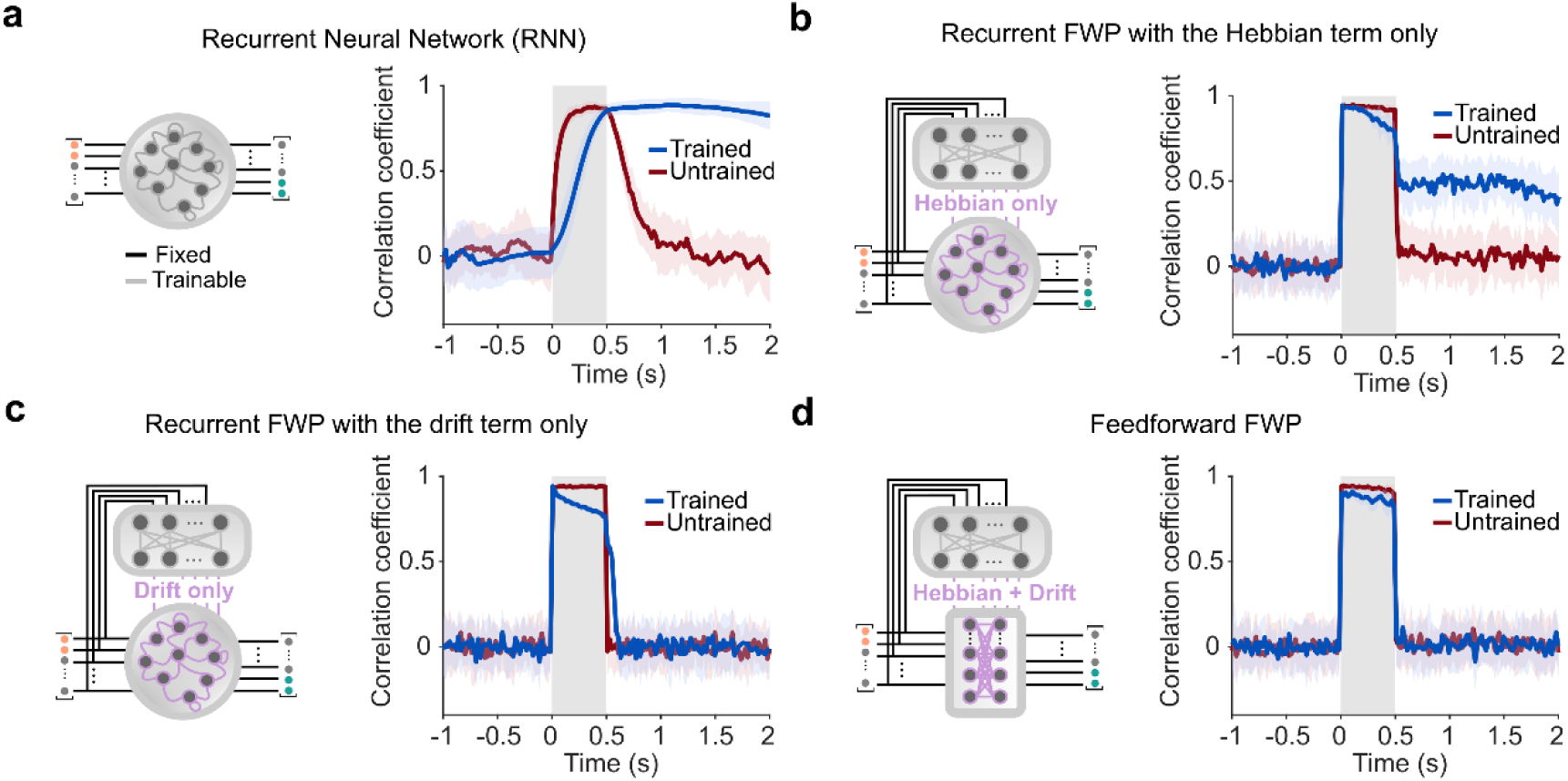
Hebbian writing, synaptic drift, and fast-net recurrence are jointly required for latent-to-mnemonic recovery. **(a)** Conventional RNN control. Left: The model had the same hidden-state architecture as the fast net, but its recurrent weights were fixed after training and received no dynamic updates from a slow net. Right: Time-resolved stimulus-neural geometry correlation in trained and untrained RNNs. **(b)** Recurrent FWP with the Hebbian component only. Left: Architecture of the ablated update rule. Right: The model maintained a stable delay-period geometry but did not show early degradation followed by late recovery. **(c)** Recurrent FWP with the drift component only. Same conventions as in (b). The model failed to maintain stimulus-specific geometry after sample offset. **(d)** Feedforward FWP. Left: Architecture of the feedforward fast net. Right: Trained and untrained models showed similar geometry trajectories, with no delay-period recovery. Gray region denotes sample presentation; light shading denotes SD across repetitions.

We next asked whether both components of the fast-weight update rule in the slow net were required. We found that removing either component disrupted the dynamics, but in different ways. When the slow net contained only the Hebbian component, the model retained a content-dependent associative update but lacked a directed synaptic drift (Fig. 5b, left; see Methods). After training, this Hebbian-only model maintained an elevated, approximately stable stimulus-geometry correlation throughout the delay (0.5–2.0 s; mean = 0.47; Fig. 5b, right), indicating that stimulus information remained continuously readable from neural activity rather than becoming latent and later reemerging (Supplementary Fig. 12). A supplementary variant using a delta-rule-style fast-weight update (Irie et al., 2021) produced the same mode of delay-period representation (Supplementary Fig. 13). Thus, associative writing alone could preserve stimulus information, but did not generate the time dependent transition from a latent state to an expressed mnemonic code.

Conversely, when the slow net contained only the drift component without a content-specific associative writing mechanism (Fig. 5c, left), the stimulus-neural geometry correlation collapsed immediately after sample offset and remained near baseline throughout the delay (0.5–2.0 s; mean = 0.03; Fig. 5c, right). Thus, the drift component alone provided a temporal evolution of the fast weights, but could not bind this evolution to the identity of the remembered stimulus.

Finally, we examined whether recurrence in the fast net were required as the original FWP uses a feedforward fast net (Schmidhuber, 1992). We trained a feedforward FWP with the same Hebbian and drift components in the slow-net update rule, but without recurrent connections in the fast net (Fig. 5d, left; see Methods). The trained and untrained feedforward FWP models showed indistinguishable stimulus-neural geometry trajectories in the delay period. Both exhibited a transient stimulus-driven geometry during sample presentation, followed by a return to baseline and no delay-period recovery (0.5–2.0 s; trained mean = 0.01; untrained mean = 0.02; 𝑡(73) = −1.36, 𝑝 = 0.18; Fig. 5d, right). Thus, even when fast weights were dynamically updated, feedforward readout was insufficient to convert the evolving synaptic state into a late-delay populational representation.

Together, these ablation results reveal a division of labor among the components of the recurrent FWP model. The Hebbian component writes stimulus-specific content into the fast weights; the drift component advances the synaptic state along a delay-period trajectory; and recurrent fast-net dynamics query the evolving synaptic state and convert it into an activity-expressed mnemonic code. Thus, the full combination of Hebbian writing, synaptic drift, and recurrent readout was required to reproduce the latent-to-mnemonic recovery observed in dlPFC.

### Fast Synaptic States Preserve Latent Memory

The preceding ablation experiments suggest that fast-weight plasticity writes stimulus-specific information into the dynamic recurrent weights. To direct examine this function, we asked whether the stimulus geometry was preserved in the synaptic state during the early delay, when the same geometry was weakly expressed in fast-net activity.

To quantify the synaptic representation, we treated the instantaneous recurrent weight matrix of the fast net as the synaptic state of the model (Fig. 6a, left). At each time point, this matrix was vectorized into a high-dimensional synaptic-state vector for each stimulus condition. We then applied the same geometry analysis used for neural activity, comparing pairwise angular relationships among stimulus-specific synaptic-state vectors with pairwise angular relationships among the stimulus locations (see Methods). We found that the stimulus-synaptic geometry correlation increased during stimulus presentation (0.05–0.25 s; mean = 0.91). Unlike the stimulus-neural geometry correlation, however, the stimulus-synaptic geometry correlation remained high throughout the delay (0.5–2.0 s; mean = 0.94), consistent with synaptic storage accounts of WM (Masse et al., 2019). Thus, during the interval in which stimulus geometry was largely absent from fast-net activity, it remained continuously represented in the dynamic fast weights.

**Figure 6:**
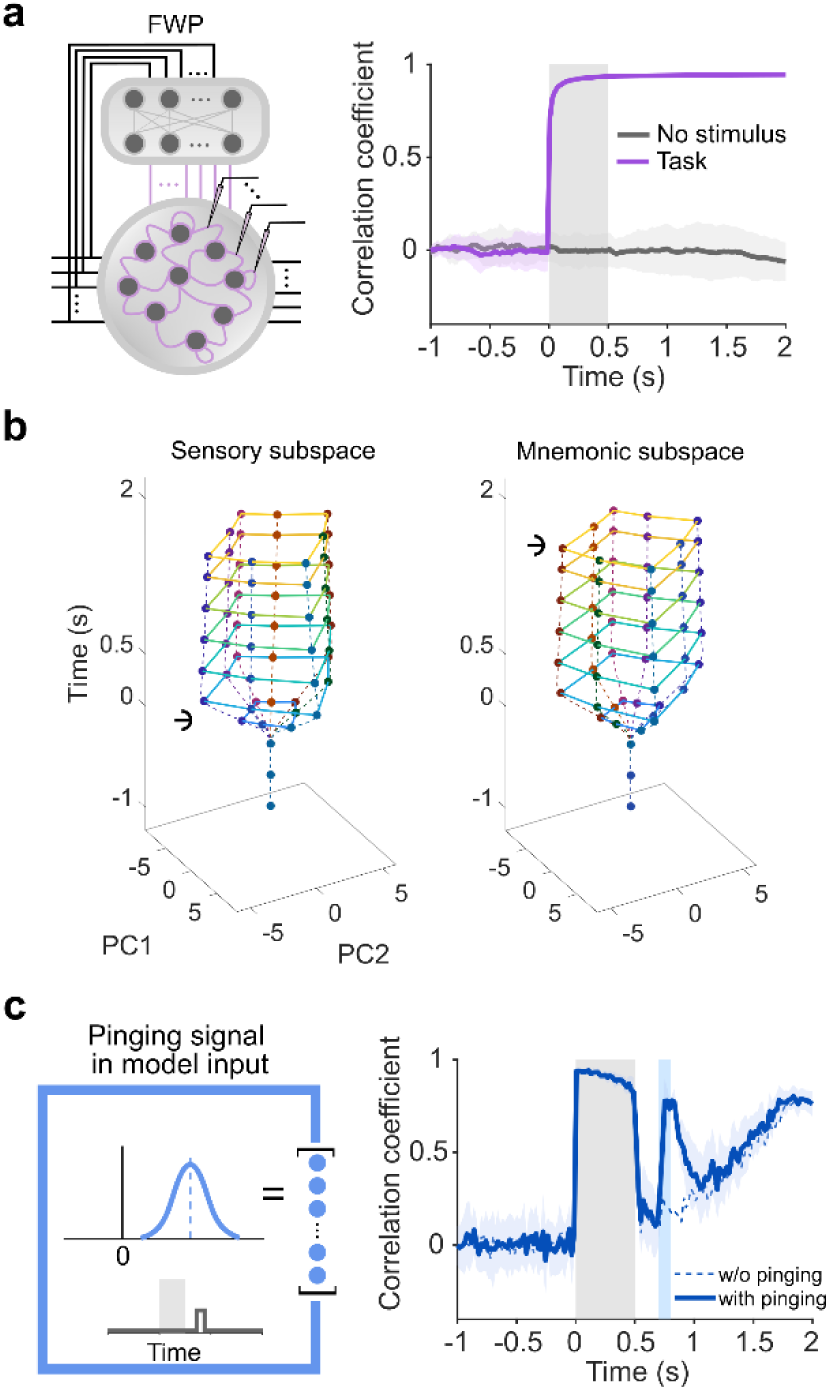
Dynamic fast weights preserve latent memory. **(a)** Synaptic geometry analysis. Left: The dynamic recurrent weights of the fast net were recorded and treated as synaptic-state variables. Right: Time-resolved stimulus-synaptic geometry correlation. Gray region denotes the sample period, and light shading denotes the SD across 20 repetitions. **(b)** Cross-temporal projections of synaptic states into the sample-period synaptic subspace (left) and the late-delay synaptic subspace (right). Anchors indicate the time point at which each subspace was defined. **(c)** Pinging 526 experiment. Left: A brief stimulus-independent Gaussian pulse was injected into all input dimensions during the early delay. Right: The perturbation transiently reinstated stimulus geometry in fast-net activity. Light shading denotes SD across 20 repetitions, blue region denotes the perturbation period, and dashed line shows the unperturbed trajectory.

Cross-temporal projection of synaptic state further revealed that this synaptic code was stable across the delay. When projected into either the sample-period synaptic subspace or the late-delay synaptic subspace, stimulus-specific synaptic states retained an ordered, stimulus-aligned geometry with relatively stable scale (Fig. 6b). The fast weights therefore did not undergo the same degradation-and-recovery seen in neural activity. Instead, they provided a latent storage medium, whereas the fast-net activity provided a time-dependent readout of that stored information.

A central prediction of the synaptic-storage account is that a nonspecific perturbation (“pinging”) delivered during an activity-silent interval should transiently reveal the latent memory (Rose et al., 2016; Wolff et al., 2017). To test this prediction in the model, we delivered a brief stimulus-independent Gaussian pulse to all input units during the early delay (Fig. 6c; see Methods), with a positive mean to simulate a global excitatory perturbation (Rose et al., 2016). This perturbation transiently reinstated the stimulus geometry in fast-net activity. The stimulus-neural geometry correlation rose abruptly from near baseline to a level comparable to the late-delay response during the perturbation window (0.7–0.8 s; max = 0.78). Cross-temporal projection further showed that the perturbation-evoked representation was expressed in the mnemonic subspace rather than a simple reinstatement in the original sensory subspace (Supplementary Fig. 14), consistent with prior empirical findings (Wolff et al., 2017). After the perturbation, the correlation returned to the unperturbed trajectory and subsequently followed the normal late-delay ramping recovery. In contrast, when the stored information in recurrent weights was erased, the ramping trajectory immediately declined to zero (Supplementary Fig. 15).

Together, these analyses show that the dynamic fast weights preserved a structured latent memory during the delay, supporting the view that fast synaptic states provide the storage medium from which mnemonic activity is subsequently reconstructed.

### Ongoing Fast-Weight Plasticity Drives Timed Memory Readout

During the late delay, the latent synaptic state was converted back into neural activity in a task-appropriate format. We first tested whether the late-delay recovery required an uninterrupted trajectory of neural activity. To do this, we clamped all hidden units of the fast net to zero at each time step during the middle delay (1.0–1.5 s), as an idealized *in silico* analogue of transient neuronal silencing (Fig. 7a, left; see Methods). This manipulation abolished the stimulus-neural geometry correlation during the inhibition window (1.0–1.5 s; mean = 0.01; Fig. 7a, right). However, immediately after release from inhibition, the correlation returned to the unperturbed trajectory and the neural states transferred to the mnemonic subspace as if no perturbation was applied (Supplementary Fig. 16), paralleling empirical observations in mouse cortex (Inagaki et al., 2019). Thus, late-delay mnemonic recovery of neural geometry did not require continuous propagation of the preceding neural state.

**Figure 7:**
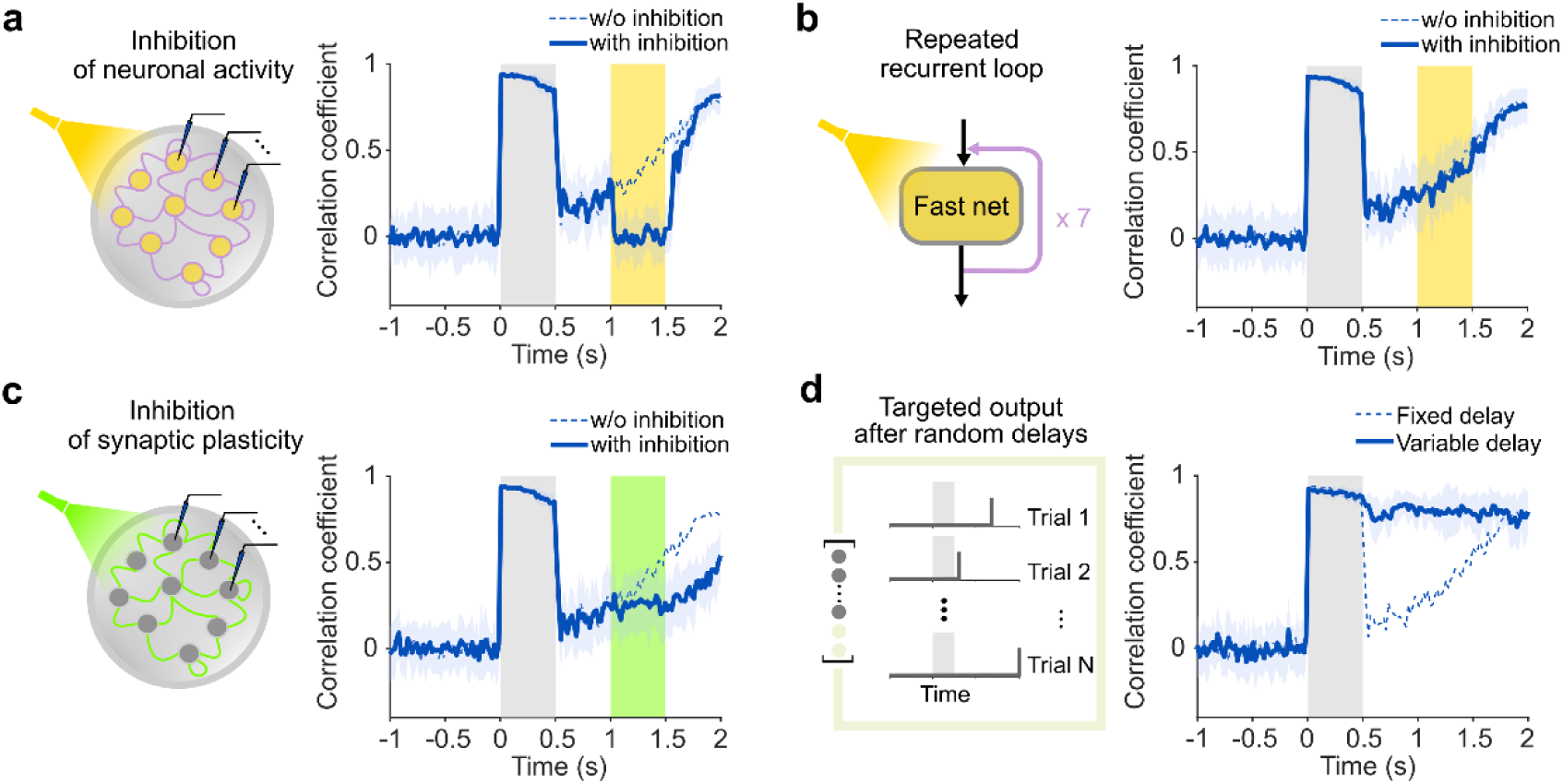
Ongoing fast-weight plasticity drives timed memory readout. **(a)** *In silico* silencing of fast-net activity. Left: All hidden activities in the fast net were clamped to zero during the middle of the delay. Right: Stimulus-neural geometry correlation with and without activity silencing. Yellow region denotes the inhibition period. **(b)** Repeated recurrent readout after activity silencing. Left: During the inhibition interval, fast-net recurrent computation was iterated seven times at each external time step. Right: Repeated recurrent computation restored the normal geometry trajectory. **(c)** Blockade of fast-weight updating. Left: The update signal from the slow net to the fast weights was set to zero during the middle delay while fast-net activity was left intact. Right: Blocking fast-weight updates attenuated the ramping recovery. Green region denotes the update-blockade period. **(d)** Variable-delay training. Left: Target outputs were imposed after randomly sampled delay lengths across trials. Right: The model trained with variable delays maintained a high stimulus-neural geometry correlation throughout the delay.

How could stimulus-aligned geometry reappear after the neural state had been erased? We hypothesized that the recurrent fast net could read out the stored synaptic state by repeatedly applying the current fast weights to nonspecific background input. To test this hypothesis, we increased recurrent computation during the same inhibition interval by iterating the fast-net recurrent loop several times at each external time step (Fig. 7b, left; see Methods). When recurrent loop was iterated enough times (seven times in this case), the stimulus-neural geometry correlation closely followed the trajectory observed without inhibition (Fig. 7b, right; see Supplementary Fig. 17 for different repetitions), although the external input during this period contained only white noise. This result indicates that the content-bearing fast weights could shape nonspecific activity into a mnemonic population pattern through recurrent readout. Therefore, recurrence in the fast net did not by itself store the memory; instead, it converted the stored synaptic state into an activity-expressed mnemonic representation.

We then examined whether ongoing fast-weight updating was required for retrieval. To do so, we blocked synaptic updating by setting the update signal from the slow net to zero during the middle delay while leaving fast-net activity intact (Fig. 7c, left; see Methods). This manipulation strongly attenuated the normal ramping recovery of stimulus geometry during the update-blockade period (Fig. 7c, right). Critically, after the blockage was released, the trajectory did not return to its unperturbed trajectory and instead remained significantly lower at the end of the delay (1.8–2.0 s; mean = 0.51; 𝑡(8) = −15.95, 𝑝 < 0.001). Importantly, the stimulus-synaptic geometry remained intact during the update-blockade period (Supplementary Fig. 18), indicating that the manipulation did not erase the stored memory content; rather, it disrupted the time-dependent synaptic evolution that made this content increasingly readable from neural activity. Thus, ongoing fast-weight plasticity had a causal role in retrieval, not merely in storage.

Further analyses indicated that this retrieval process was primarily driven by the drift component of the fast-weight update rule (Supplementary Fig. 19). In the fixed-delay condition, the output was required only at a predictable time, so the learned drift term could act as a synaptic timing mechanism, gradually advancing the fast-weight state toward a configuration that reconstructed the stimulus geometry near the end of the delay. The ramp trajectory therefore emerged as a learned solution for timed retrieval (Durstewitz, 2004). This interpretation predicts that when the model cannot predict when memory will be required, it should keep the mnemonic representation continuously available rather than delay retrieval until a fixed endpoint (Stroud et al., 2024). To test this prediction, we trained a recurrent FWP model in which the target outputs were imposed after variable delays, uniformly sampled between 0 and 1.5 s across trials (Fig. 7d, left; see Methods). Under temporal uncertainty, the model no longer showed the early-delay degradation followed by late-delay recovery observed in the fixed-delay model. Instead, the stimulus-neural geometry correlation remained high throughout the delay (0.5–2.0 s; mean = 0.79, Fig. 7d, right), and cross-temporal analysis showed a more stable delay-period code (Supplementary Fig. 20), consistent with empirical observations (Inagaki et al., 2019). Therefore, the same fast-weight architecture could express memory either as a latent-to-active ramping trajectory or as a continuous available mnemonic code, depending on when the task required retrieval.

Together, these perturbation and variable-delay analyses identify fast-weight plasticity as the mechanism controlling mnemonic readout. This mechanism explains how memory can be maintained silently, recovered after transient activity silencing, and retrieved either gradually or continuously depending on temporal task demands.

## Discussion

In this study, we propose that biological working memory can be interpreted through an algorithmic principle that is shared with the Transformer family (Supplementary Fig. 21). Information is not simply held in persistent neural activity, but is written into a rapidly modifiable weight state and later queried to generate task-relevant neural activity. Combining neural-geometry analyses of macaque dlPFC recordings with a recurrent fast-weight programmer model, we show that WM can be understood as recurrent fast-weight computation. In this process, transient sensory activity programs a synaptic-like state that preserves latent information, and recurrent circuit dynamics later read from this state to express the remembered content in a partially distinct mnemonic subspace. This framework links an activity-silent and ramping-retrieval WM, a central phenomenon in neuroscience, to fast-weight and linear-Transformer computation, a central algorithmic family in modern AI. It therefore suggests that biological and artificial systems may converge on a common solution for programmable temporary memory.

A central advance of our study is to extend synaptic accounts of WM from latent maintenance to active retrieval. Previous synaptic theories have shown how memory can be maintained silently (Mongillo et al., 2008; Mi et al., 2017; Panichello et al., 2024; Mongillo & Tsodyks, 2026), but they have not fully specified how latent content becomes reinstated in neural activity. The recurrent FWP model provides a mechanistic account of this transition. Hebbian writing stores stimulus-specific content into fast weights, synaptic drift advances this state over time, and recurrent circuit dynamics query the evolving synaptic state to express the memory in a task-appropriate format. From this perspective, ramping activity needs not reflect a slowly accumulating firing-rate signal or an external timing input; instead, it reflects the visible neural readout of an evolving synaptic memory. This interpretation also clarifies perturbation-evoked reactivation. In the model, a nonspecific pulse probes a circuit whose fast weights already contain the latent memory, therefore transiently converting hidden synaptic structure into neural activity. Pinging therefore becomes a computational query to a latent state. Importantly, the reactivated representation emerges in the mnemonic subspace rather than as a replay of the original sensory response, explaining how a WM representation can be silent, recoverable, and already formatted for future behavior.

The present findings also establish a direct computational link between biological WM and the Transformer family through fast-weight computation. Linear Transformers have been shown to be equivalent to FWPs (Schlag et al., 2021), in which key-values outer products write information into a dynamic matrix-valued memory and query vectors read from this memory during sequence processing. From a WM perspective, this dynamic matrix is not simply an attention device, but a transient memory buffer (Gershman et al., 2025). The outer product update resembles Hebbian writing (Irie & Gershman, 2025), while the query operation resembles retrieval from a latent synaptic state. Thus, linear-Transformer computation can be interpreted as a cycle in which information is written into, organized within, and queried from a dynamically changing memory state. On the other hand, biological WM can be interpreted as a recurrent fast-weight system in which neural activity programs synaptic states and recurrent circuit dynamics query these states to guide behavior. This reciprocal interpretation gives Transformer architectures a biological grounded computational meaning, while giving synaptic theories of WM a mathematically explicit language for describing storage and retrieval.

This equivalence should be understood as algorithmic rather than literal. Biological WM is not a standard Transformer, nor do we assume that cortical circuits implement attention in the engineering form used in modern artificial systems. The relevant point is more general. Both systems use dynamically updated weights as an internal memory state that can be written, organized, and queried. This perspective suggests that the success of Transformer-family architectures may not depend solely on attention conceived as pairwise token matching (Vaswani et al., 2017), but may also reflect a broader principle of programmable temporary memory. Furthermore, biological WM may offer additional constraints for this principle, because it operates under severe capacity limits (Mi et al., 2017) while replying on chunking (Nassar et al., 2018), gating (Chatham & Badre, 2015), and hierarchical organization (Fan et al., 2025) to support flexible cognition. For example, recent analyses of linear-attention models suggest that compressed memory matrices can develop inefficient or redundant internal structures as sequence information accumulates (Sun et al., 2026). In this context, a carrier dynamic such as the slow synaptic drift in our model may provide a useful organizing mechanism that reshapes the geometry of memory states over time without overwriting stored content, allowing temporary memory to remain neither inert nor unstable, but dynamically prepared for retrieval.

Recurrence is central to this account. Standard linear Transformers read from the fast-weight state through a largely feedforward query operation, whereas cortical circuits supporting WM are intrinsically recurrent. Our results suggest that recurrence is more than a biological details appended to Transformer-family computation. It is the operation that converts latent synaptic states into an expressed neural representation. In the recurrent FWP model, fast weights provide a temporary matrix-valued memory, and recurrent dynamics repeatedly query this memory over internal time. This iterative readout allows stored content to be extracted, transformed, and expressed in a behaviorally-appropriate mnemonic subspace. The model therefore extends the linear Transformer view of fast-weight memory into a circuit level mechanism for memory retrieval. This extension is consistent with the broader neuroscience-inspired AI program, in which biological principles can help identify algorithmic constraints for more flexible artificial systems (Hassabis et al., 2017). In this regard, looped Transformer architectures (Saunshi et al., 2025) are relevant because they suggest that repeated computation over internal representations can improve reasoning performance.

Together, these findings recast WM as a programmable fast-weight process, rather than as a choice between persistent neural activity and silent storage. In this process, sensory activity writes information into rapidly modifiable synaptic states, synaptic dynamics organize this content over time, and recurrent circuits query the evolving state to generate behaviorally-relevant neural activity. This framework unifies activity-silent maintenance, timed mnemonic retrieval, and Transformer-family computation within a single algorithmic account of temporal memory in biological and artificial intelligence. Several important limitations remain. The empirical analyses measured neural activity rather than synaptic states directly, so the proposed synaptic mechanism is inferred through modelling rather than observed *in vivo*. The present model also focused on memory writing and readout, leaving the mechanisms that clear or overwrite fast-weight memory for future work. In addition, the link to Transformers is computational, not anatomical or implementational.

Nevertheless, the success of Transformer-family architectures across diverse AI tasks raises the possibility that fast-weight synaptic principles may extend beyond WM to other cognitive domains. Determining whether latent to active recovery, synaptic-state organization, and plasticity-mediated readout occur across cognitive domains will test whether programmable fast-weight computation is a broader algorithmic principle shared by biological and artificial intelligence.

## Acknowledgments

We thank Drs. Christos Constantinidis and Hua Tang for guidance on preprocessing the neurophysiological data, and Mr. Tao Liu for valuable discussions and comments.

## Author contributions

Conceptualization: L.J. and J.L. Methodology: L.J., Y.Z., and J.L. Data curation: L.J. Validation: L.J., Y.Z., and J.L. Software: L.J. Investigation: L.J. and Y.Z. Formal analysis: L.J. and Y.Z. Visualization: L.J. Resources: J.L. Supervision: J.L. Project administration: J.L. Funding acquisition: J.L. Writing—original draft: L.J. and J.L. Writing—review and editing:

## Methods

### Experimental design

We reanalyzed neurophysiological data from three rhesus macaques (*Macaca mulatta*; monkeys ELV, ADR and NIN) performing a visuospatial delayed-match-to-sample task, originally reported in previous studies (Meyer et al., 2011; Qi et al., 2011; Tang et al., 2022). Monkeys were seated in a primate chair while a monitor was placed 60 cm away. They fixated on a 0.2° white square while a 2° white square appeared pseudo-randomly at one of nine locations arranged in a 3×3 grid with 10° spacing in between, including the central fixation location. Two experimental conditions were compared: passive viewing and active decision-making. In the passive condition, monkeys received liquid rewards for maintaining fixation; in the active condition, rewards were contingent on executing saccades to correct targets. Target locations were orthogonal to stimulus locations and varied randomly across trials, preventing advance saccade planning. Error trials were infrequent (8% error rate; Kobak et al., 2016) and only correct trials were included in the analysis. The monkeys were familiarized with all the stimuli before the recorded experiment sessions.

### Neural recordings

Recording sites were located in the dorsolateral prefrontal cortex, encompassing the principal sulcus and extending posterior to the arcuate sulcus, including area 46 and portions of area 8a. Details regarding surgical procedures, recording equipment and preprocessing that converted physiological signals into spike trains are provided in (Meyer et al., 2011; Qi et al., 2011; Tang et al., 2022). To compute peristimulus time histograms (PSTHs), spike trains were convolved with a Gaussian kernel (SD = 10 ms) and sampled at 500 Hz. The resulting PSTHs were truncated to the working memory period and averaged across all trials, as well as separately across match and nonmatch decision trials. These trial-averaged PSTHs served as each neuron’s firing rates in response to stimuli. The number of recorded neurons varied across monkeys and conditions. Monkey ELV yielded the largest number of recordings, which were balanced across the two conditions (384 neurons per condition). Accordingly, results from monkey ELV are presented in the main text, whereas those from monkeys ADR and NIN are provided in the Supplementary Figs. 1 and 6.

### Linear decoding

To classify neural population representations of diametrically opposite stimuli, we selected neurons with ≥16 experimental trials per stimulus condition (226 neurons total). For each neuron, trials were randomly split into two equal-sized subsets for training and testing, respectively. Mean firing rates were computed for each subset, yielding 226-dimensional neural states representing each stimulus condition. Time-resolved decoding performance was quantified using a 200-ms sliding window (100 time points per window) advanced in 20-ms steps. A linear support vector machine (SVM) was trained and tested on the neural states to obtain decoding accuracy for each time window. To estimate variability due to trial assignment, the random split procedure was repeated 10 times.

### Cross-temporal decoding

To assess whether decoders trained during the sample period would generalize to other task epochs, we trained linear SVM decoders to discriminate between all pairwise combinations of the eight stimulus conditions (28 decoders total). Training used trial-averaged firing rates from 0.05 to 0.45 s (200 time points per window). The trained decoders were then tested on the firing rates from the fixation (−0.95 to -0.55 s and -0.45 to -0.05 s), the early delay (0.55–0.95 s), the middle delay (1.05–1.45 s), late delay (1.55–1.95 s), and the probe periods (2.05–2.45 s).

For a more comprehensive cross-temporal decoding analysis, we applied a 200-ms sliding window advanced in 20-ms steps across the sample and delay periods. The 28 decoders trained at each time window were tested on all windows. Mean decoding accuracies for each testing window within the delay period were compiled into a 70×70 decoding accuracy matrix. The diagonal region of this matrix, defined with a margin of 20 cells on each side, separated the upper-left and lower-right triangular regions.

### Stimulus-neural geometry correlation

Although most of the 384 neurons of monkey ELV were recorded individually, they were analyzed as a pseudo-population, as if recorded simultaneously. Population activity for the nine stimulus conditions within a 50-ms time window formed nine clusters in the 384-dimensional neural state space. Structure of the cluster centroids defined a geometry. This geometry was characterized by aRDVs: Vectors were draw from the cluster centroid of the fixation location to that of the eight stimulus locations; angles between any two vectors were recorded in an aRDV which had 28 elements in total. This aRDV was z-scored and concatenated with three z-scored aRDVs in consecutive time windows to create a 112-element grand aRDV for a 200-ms time window (Supplementary Fig. 2). The neural grand aRDV was compared with the grand aRDV of the stimulus geometry, which was created by drawing vectors from the central fixation location to the eight stimulus locations in a 2-dimensional space, calculating the aRDV, z-scoring, and concatenating the aRDV with three self-copies. The time-resolved Pearson’s correlation coefficient was computed using a 200-ms wide sliding window which advanced in 20-ms steps. The baseline was established by randomly permuting the stimulus labels 100 times prior to computing the correlation.

### PCA visualization

We visualized dimension-reduced neural geometry in three 200-ms wide time windows: 0.05–0.25 s, 0.6–0.8 s, and 1.8–2.0 s. PCA was applied to cluster centroids of the eight stimulus conditions in the neural state space during each time window. The high-dimensional cluster centroids were projected to the two PCs with the most explained variances for visualization.

### Cross-temporal PCA projection

To project high-dimensional neural states onto the 2-dimensional sensory subspace, we applied PCA to the neural activity within a 0.2 s window of the sample period, using the top two principal components (PCs) as the subspace basis. Neural states from all other time windows were then projected onto this basis. Similarly, the mnemonic subspace basis was defined by the top two PCs obtained during a 1.9 s window in the late delay period. To compare the scale of the neural geometries between these two epochs, we quantified the area enclosed by the projected neural states using the shoelace formula.

### Representation transfer task

We trained computational models to perform the representation transfer task. The models processed input sequences spanning 150 timesteps, corresponding to 3-s trial length in the animal experiment. Each sequence was divided into three distinct periods: a 50-step fixation period, a 25-step sample period, and a 75-step delay period. Spatial location cues were presented during the sample period as 2-dimensional coordinates sampled from a 3×3 grid, defined as follows,

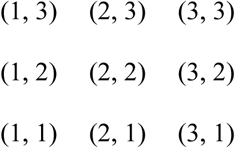

where (2, 2) served as the reference location, while the remaining eight coordinates functioned as stimulus locations. During sample presentation, these coordinates were embedded in the first two dimensions of a 384-dimensional input vector, with all remaining elements set to zero. During the fixation and delay periods, all elements of the input vector were set to zero. To simulate experimental noise, independent and identical distributed zero-mean Gaussian noise with standard deviation 0.1 was added to each dimension of the input vector at each timestep.

The models generated outputs exclusively at the final timestep. The target outputs were 384-dimensional vectors containing the location cues in their final two dimensions, with all other elements set to zero. Thus, the objective of the task was to transfer location cues from the first two dimensions of the input vectors to the last two dimensions of the output vectors.

### Recurrent FWP model

The recurrent FWP model consisted of a feedforward slow net and a recurrent fast net. Both the two nets accept the input sequences [𝑥_1_, 𝑥_2_, … , 𝑥_150_]. For input vector 𝑥_𝑡_, the slow net computed the key vector 𝑘_𝑡_ and value vector 𝑣_𝑡_ through the mapping

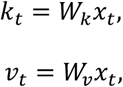

where 𝑊_𝑘_ and 𝑊_𝑣_ were trainable matrices which were initialized randomly. The key and value vectors constituted the Hebbian-like associative component in the fast-weight update rule through outer product 𝑣_𝑡_𝑘^𝑇^. A trainable constant matrix 𝐷, which was initialized randomly, constituted the drift component. The fast-weight update signal at each time step was

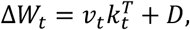

Additionally, in the fast-weight update ablation analysis, the Hebbian-only update signal was set as

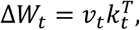

and the drift only update signal was set as

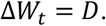

The update signal modified the recurrent weights of the fast net at each time step as follows,

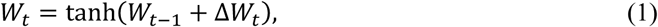

where tanh(⋅) was the squashing function for limiting the weights in a finite range and 𝑊_0_ = 0. We also trained a recurrent FWP model with linear weight updating, that is 𝑊_𝑡_ = 𝑊_𝑡−1_ + Δ𝑊_𝑡_. The results were similar to those obtained from the model using tanh(⋅) (Supplementary Fig. 22).

The fast net has 384 hidden units. The hidden state ℎ_𝑡_ was updated recurrently by

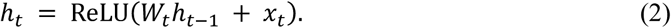

where ReLU(⋅) was the rectified linear unit function and ℎ_0_ = 0. The last hidden state was the model output: 𝑦 = ℎ_150_.

### Recurrent FWP training and testing

We used a supervised learning paradigm with training pairs consisting of input sequences and target outputs. The input sequences combined spatial location cues with random Gaussian noise. To construct the training set, input sequences corresponding to each of the nine location cues were sampled 8 times, yielding a total of 72 data pairs. Training minimized a masked mean-squared error loss applied to the model outputs. The mask was restricted exclusively to the first and last two dimensions of the output vectors, excluding all other dimensions from the objective function. A recurrent FWP model was trained using the Adam optimizer using a learning rate of 1 × 10^−6^ and a batch size of 9 for 1000 epochs. Model parameters yielding the minimum epoch-averaged loss were saved for analysis.

In addition to the fixed-delay task, a separate recurrent FWP model was trained on a representation transfer task featuring variable delay lengths. During training, the model output was evaluated against its target after a delay duration uniformly sampled between 1 to 75 timesteps for each batch. To ensure sufficient coverage of this variable delay range, each location cue was sampled 40 times to create 360 training data pairs. All other hyperparameters and training configurations remained identical to the fixed-delay task.

Following training, model performance was evaluated across 20 independent test trials. For each trial, a novel input sequence was independently sampled and fed into the recurrent FWP models, and the resulting outputs were compared against their target outputs. Hidden unit activations and the recurrent weight matrices of the fast net were recorded throughout these test trials for further analysis.

### Fast-net activity analysis

We analyzed the hidden unit activity of the fast net using the same analytical pipeline applied to the dlPFC recordings. For cross-temporal PCA projection, a 10-step sliding window with a 1-step stride was applied to the trial-averaged activity. The sensory and mnemonic subspaces were then constructed using the activity at the 60th and 145th timesteps, respectively. For cross-temporal decoding, linear SVMs were trained to decode 28 unique pairs of peripheral sample locations in a 10-step sliding window with a 1-step stride. The mean accuracies across all 28 decoders were compiled to form the decoding accuracy matrix. For stimulus-neural geometry correlation, neural and stimulus aRDVs were constructed at each timestep, and their Pearson’s correlation coefficients were computed for each testing trial. Finally, two-dimensional PCA projections capturing the sample presentation, early delay, and late delay periods were applied to a representative test trial.

### Recurrent neural network

The baseline RNN cell contained 384 hidden units, accepted a 150- timestep, 384-dimensional input sequence, and generated a 384-dimensional output exclusively at the final timestep. The hidden state ℎ_𝑡_ was recurrently updated by ℎ_𝑡_ = tanh(𝑊ℎ_𝑡−1_ + 𝑥_𝑡_), where 𝑊 represents the trainable recurrent weight, initialized as an orthogonal matrix. The output was generated according to 𝑦 = ℎ_150_.

The training data were sampled identically to the recurrent FWP model, with the exception that the spatial coordinates of the location cues were shifted by subtracting two. This adjustment accommodated the [-1, 1] dynamic range of the tanh(⋅) activation function. Each location cue was sampled 80 times to generate 720 training data pairs. The model was trained using the Adam optimizer using a learning rate of 1 × 10^−6^ and a batch size of 9 for 1000 epochs. Model evaluation was conducted across 20 trials using independently sampled, novel data.

### Feedforward FWP

The feedforward FWP model utilized the same slow net configuration, fast-weight update rule, and training and testing procedures as the recurrent FWP model. However, the hidden state ℎ_𝑡_ in the fast net was updated without recurrent feedback: ℎ_𝑡_ = ReLU(𝑊_𝑡_𝑥_𝑡_ + 𝑥_𝑡_), where 𝑊_𝑡_ and 𝑥_𝑡_ represent the dynamic fast weights and the current input, respectively; and initial values of ℎ_𝑡_ and 𝑊_𝑡_ were set to ℎ_0_ = 0 and 𝑊_0_ = 0. The model output was generated via 𝑦 = ℎ_150_.

### Synaptic state analysis

During testing of the recurrent FWP model, the 384×384 recurrent weight matrix 𝑊_𝑡_ was recorded at each timestep. To define the dynamic synaptic state, this matrix was flattened into a 147456-dimensional vector 𝑠_𝑡_. The synaptic geometry was then constructed using the synaptic states corresponding to each of the location cues. To evaluate the structural properties of these configurations, stimulus-synaptic geometry correlation and cross-temporal projection were computed using the identical analytical pipeline applied to the empirical data. As a control, a “no-stimulus” condition was implemented in which no location cue was embedded in the input vectors during sample presentation.

### Pinging simulation

To simulate the application of an uninformative “pinging” stimulus, we injected positive-mean Gaussian noise into all input dimensions during a designated pinging period. Specifically, during model testing, input sequences were sampled as described above, with the exception that between the 85th and 90th timesteps, the mean of the Gaussian noise was shifted to 0.2 with the standard deviation remaining at 0.1. Crucially, this positive-mean pulse carried no spatial information regarding the location cues, serving as a purely non-specific input to probe the latent states of the network.

### Neuronal activity inhibition

We simulated optogenetic inhibition of neuronal activity by temporarily suppressing hidden state activity in the fast net of the recurrent FWP model. Specifically, while the recurrent weights 𝑊_𝑡_ and the internal hidden state ℎ_𝑡_ in the fast net updated normally according to Eqns. (1) and (2), the post-update hidden state ℎ_𝑡_ at each 𝑡 between the 100th and 125th timesteps was overwritten. During this inhibition period, ℎ_𝑡_ was replaced across all dimensions by zero-mean Gaussian noise with standard deviation of 1 × 10^−10^. Consequently, stimulus-related information was completely abolished from the active neural state ℎ_𝑡_ during the period, allowing evaluation of the specific contribution of neuronal activity.

### Repeated recurrent computation

During normal testing, recurrent computation in the recurrent FWP model updated exactly once every timestep. We relaxed this constraint by allowing multiple recurrent updating at each timestep during the neuronal activity inhibition period. Specifically, at a timestep 𝑡 between the 100th and 125th timesteps, the hidden state ℎ_𝑡_ was initially clamped to zero. Then, the following iterative equation was applied

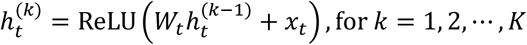

Where 𝑥_𝑡_ represents the input vector at the current timestep, K denotes the total number of recurrent update steps, and the initial condition ℎ_t_^(0)^ = 0. For timesteps outside this inhibition period, the model proceeded with its standard, single-step update execution.

### Inhibition of synaptic plasticity

We simulated optogenetic inhibition of synaptic plasticity within the recurrent FWP model by temporarily blocking the recurrent weight updating. Specifically, during model testing, at each 𝑡 between the 100th and 125th timesteps, fast-weight update signal Δ𝑊_𝑡_ was set to zero. Consequently, the fast-weight matrix remained constant throughout this period, preventing any activity-dependent synaptic modifications. For timesteps outside this inhibition period, the model proceeded with its standard weight-update protocol.

