## Supplementary Information for "Working Memory as Programmable Fast Weight Computation"

In Supplementary Material, we present 22 additional figures to further explain the methods and support the results in the main text.

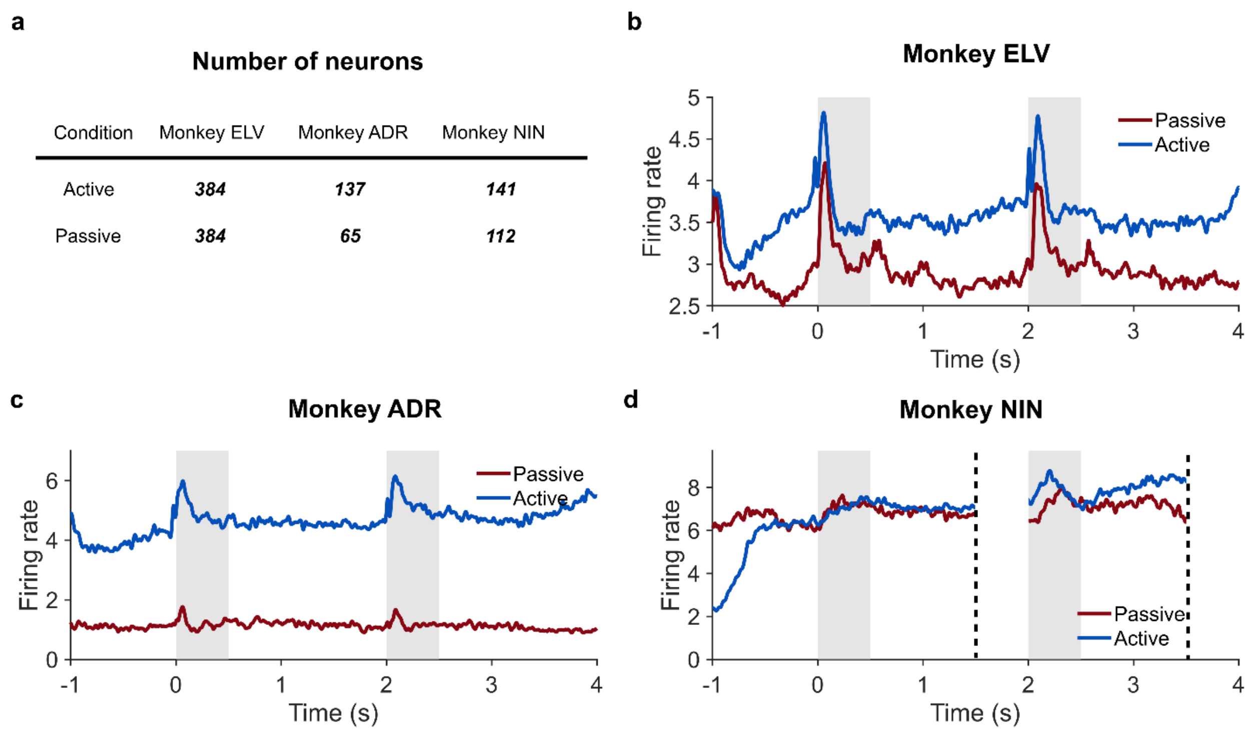


**Figure 1. Recorded neuron counts and averaged peri-stimulus time histograms.** **(a)** Number of recorded neurons in the dorsolateral prefrontal cortex (PFC) of each monkey under the active and passive conditions. In total, 384 neurons from monkey ELV were recorded across both conditions. For monkey ADR, 137 neurons were recorded in the active condition and 65 in the passive condition. All recorded neurons were included without further selection. For monkey NIN, because each neuron was recorded in tasks employing variable delay lengths (0.35–1.5 s), only neurons with delay lengths exceeding 1 s were selected, and their delay periods were truncated to 1 s before being assembled as a pseudo-population. The number of selected neurons for monkey NIN was 141 in the active condition and 112 in the passive condition. **(b)** Peri-stimulus time histogram averaged across all recorded neurons from monkey ELV in both tasks. The gray region indicates the sample and probe presentation periods. The population average reveals clear responses to the sample and probe, with the mean firing rate in the passive condition lower than that in the active condition. **(c)** Same as (b), but for monkey ADR. **(d)** Same as (b), but for monkey NIN. Both delay periods were truncated to 1 s.

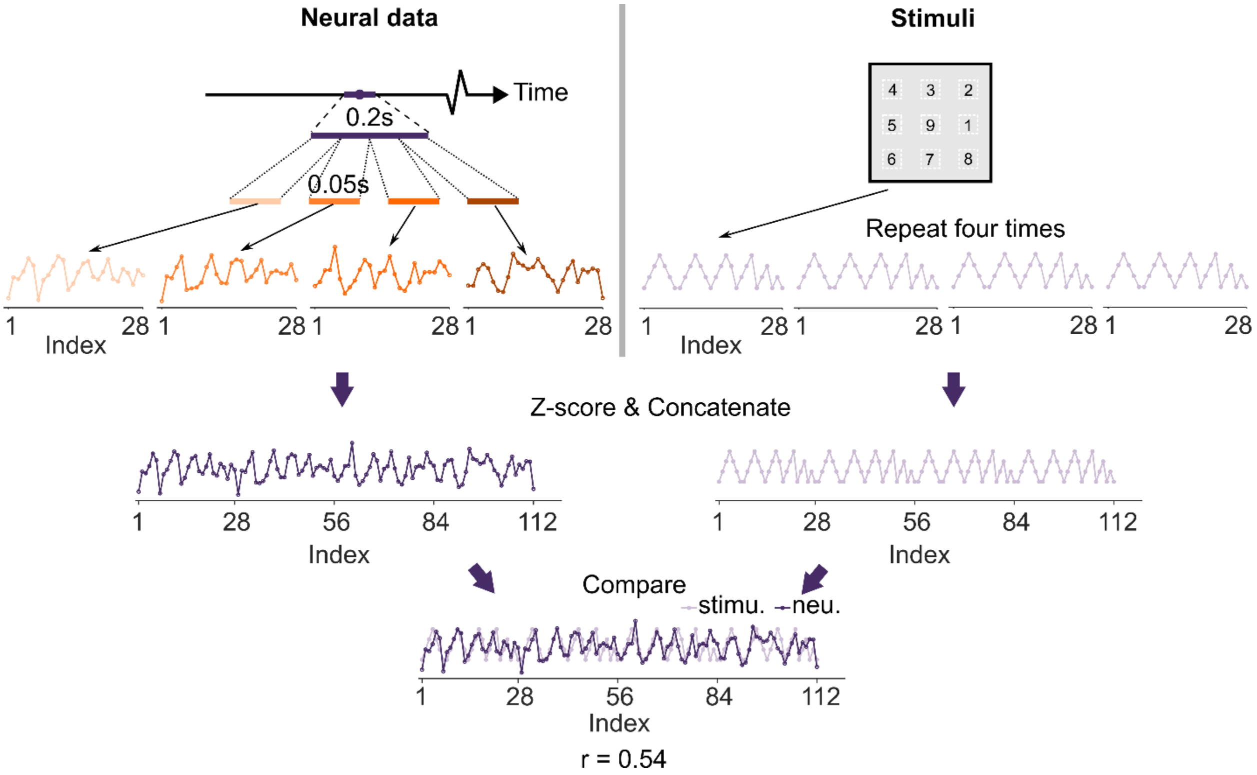


**Figure 2. Construction of the angular representational dissimilarity vector (aRDV).** Left: To compute the neural aRDV, we applied a sliding time window of 0.2 s width with a step size of 0.02 s. To enhance correlational robustness, the sampling window was partitioned into four equal sub-windows. Within each sub-window, a 28-component aRDV characterizing the neural geometry was derived. The aRDVs were then z-scored to remove spurious correlations arising from magnitude differences across sub-windows, and the resulting vectors were concatenated into a single 112-component pattern. Right: The 28-component RDV of the stimulus geometry was replicated four times, z-scored, and concatenated to form a matching 112-component pattern. Pearson's correlation coefficient was then computed between the two concatenated patterns. This procedure increased measurement stability when the patterns were uncorrelated while preserving a faithful representation of genuine geometric similarity. The baseline distribution was constructed by randomly permuting the stimulus labels 100 times and computing the correlation of the resulting permuted stimulus geometries.

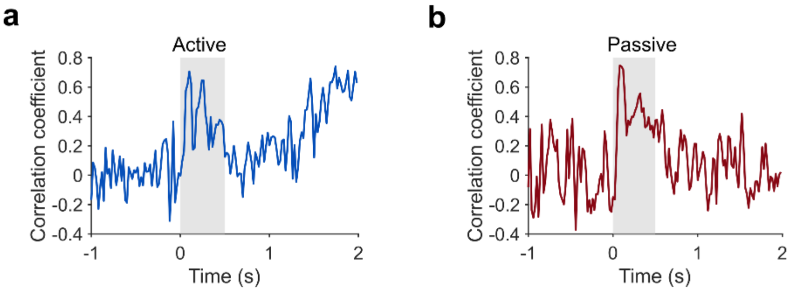


**Figure 3. Late delay period stimulus–neural geometry correlation was comparable to that during sample presentation in the absence of windowed smoothing.**

In the main text, the stimulus–neural correlation in the late delay period of the active condition appears higher than the correlation during sample presentation. This elevation is an artifact introduced by the sliding window used for smoothing. Here, we present the time-resolved stimulus–neural correlation computed without smoothing. **(a)** Time-resolved stimulus–neural correlation in the active condition. During the delay, the correlation increased to a level where the peak value (1.8–2.0 s, maximum = 0.71) matched the peak correlation observed during sample presentation (0.05–0.25 s, maximum = 0.71). **(b)** Time-resolved stimulus–neural correlation in the passive condition. The peak correlation during sample presentation was slightly higher than in the active condition (0.05–0.25 s, maximum = 0.75).

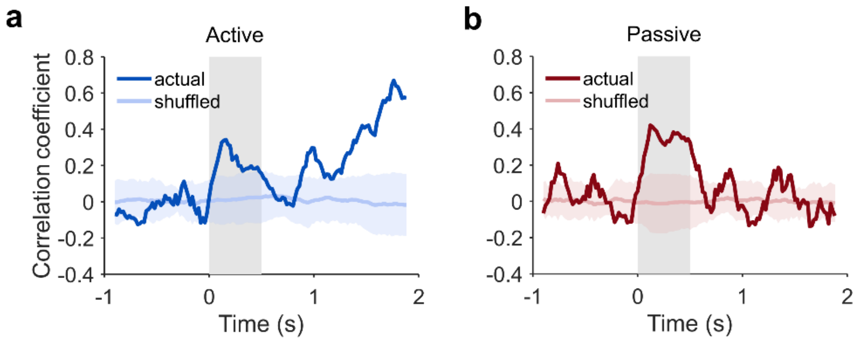


**Figure 4. The degradation-and-recovery pattern in the active condition was also evident when using distance-based RDVs.**
In the main text, stimulus–neural geometry similarity is quantified using angular RDVs (aRDVs). As an alternative, we employed distance RDVs (dRDVs). Specifically, for the stimulus geometry comprising eight locations, we computed the Euclidean distances between every pair of stimuli and assembled these distances into a dRDV vector. A dRDV was similarly constructed for the neural geometry, and the correlation between the stimulus and neural dRDVs was computed. **(a)** Time-resolved Pearson's correlation between the stimulus geometry dRDV and the neural geometry dRDV across the trial in the active condition. The gray region denotes sample presentation. The baseline was established by randomly shuffling the stimulus labels 100 times. The mean (light-colored line) and standard deviation (light shaded area) are displayed. During the delay period, the correlation first decreased and then increased, mirroring the pattern observed with aRDVs. **(b)** Time-resolved correlation in the passive condition. The correlation decreased and fluctuated near zero, consistent with the results obtained using aRDVs.

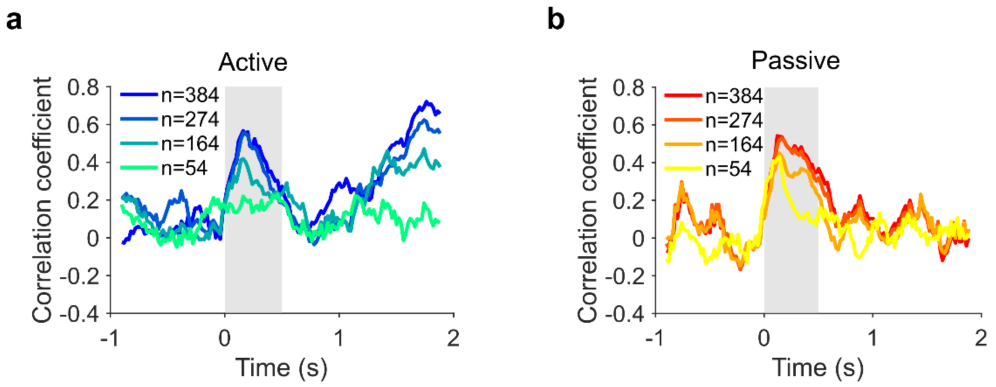


**Figure 5. Randomly reducing the size of the neural population attenuates ramping activity in the late delay period.**

We randomly removed neurons from the population and computed the stimulus–neural geometry correlation for each size-reduced population. **(a)** Time-resolved stimulus–neural geometry correlation for neural populations of varying sizes in the active condition. Smaller populations exhibited reduced ramping activity during the late delay period. **(b)** Same as (a), but for the passive condition. The difference among the populations was less prominent than in the active condition.

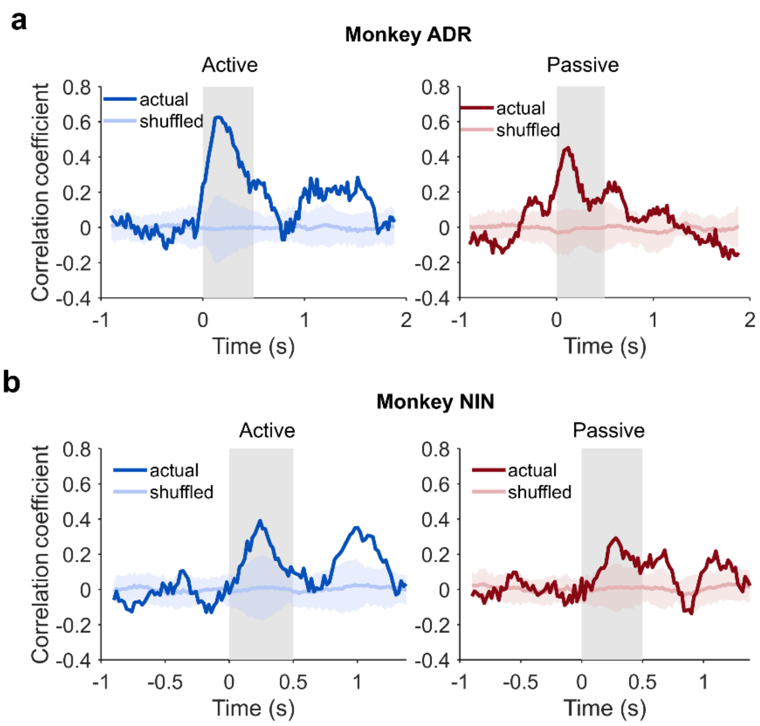


**Figure 6. The degradation-and-recovery pattern was observed in the other two monkeys.** In the main text, the stimulus–neural geometry correlation was computed from neural recordings in monkey ELV, as this subject provided the largest number of recorded neurons in both the active (384 neurons) and passive (384 neurons) conditions. The remaining two monkeys, ADR and NIN, yielded smaller sample sizes. We computed the time-resolved stimulus–neural geometry correlation for both monkeys in each condition. **(a)** Left: Time-resolved stimulus–neural geometry correlation derived from recordings in monkey ADR during the active condition. Although the correlation did not exhibit ramping activity in the late delay period—potentially attributable to an insufficient number of recorded neurons (Supplementary Fig. 5a)—it degraded during the early delay and reemerged in the middle delay, a pattern consistent with the results from monkey ELV. Right: Time-resolved correlation in the passive task. **(b)** Left: Time-resolved correlation for monkey NIN in the active condition. The degradation-and-reemergence pattern was evident, although continuous ramping was absent. Right: Time-resolved correlation in the passive task. Gray regions denote the sample presentation period. Baselines were generated by permuting stimulus labels 100 times.

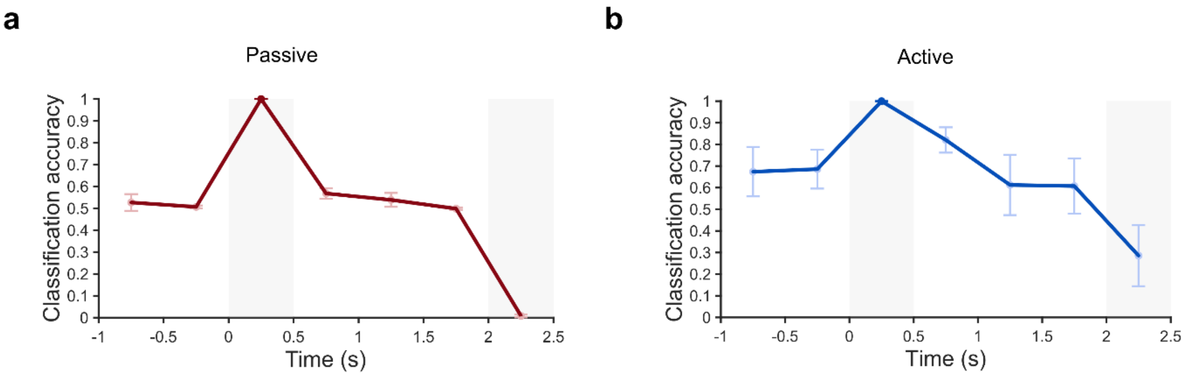


**Figure 7. Testing accuracies in the probe period of the decoders trained in the sample period were inverted in nonmatch trials.**

In the main text, cross-temporal decoding of stimulus location was performed using trials in which the sample and probe matched. Here, we conducted an analogous analysis on nonmatch trials, in which the probe appeared at the location diametrically opposite to the sample. A linear support vector machine (SVM) was trained to discriminate diametrically opposite stimuli within the 0.05–0.45 s window. The decoders were subsequently tested across the following task epochs: fixation (−0.95 to −0.55 s and −0.45 to −0.05 s), early delay (0.55–0.95 s), middle delay (1.05–1.45 s), late delay (1.55–1.95 s), and probe (2.05–2.45 s). **(a)** Cross-temporal decoding in the passive condition. Decoding accuracy during probe presentation was zero, corresponding to perfectly inverted decoding. **(b)** Cross-temporal decoding in the active condition. The mean accuracy during the probe period was 0.29, which also fell below the chance level of 0.5.

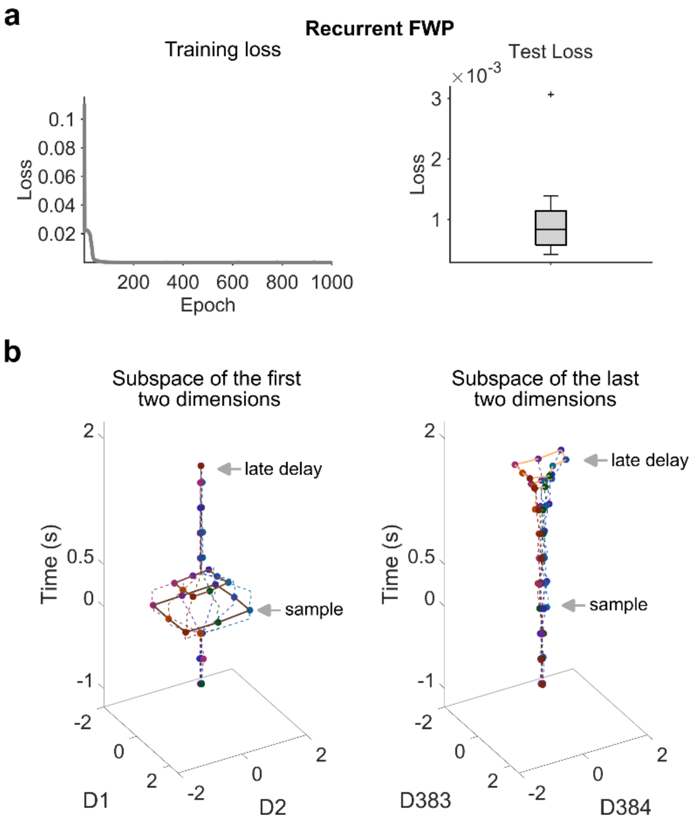


**Figure 8. A recurrent FWP model was successfully trained to execute the representation transfer task.**

**(a)** Left: Over the course of 1,000 training epochs, the loss decreased and converged to near zero. Right: When evaluated on additionally sampled test data, the test loss remained at a low level (median = 0.0008). **(b)** Left: The hidden states across the trial were projected onto the subspace spanned by the first two dimensions. Stimulus-aligned neural geometry emerged transiently in this subspace during sample presentation but was absent following sample offset. Right: The hidden states were projected onto the subspace spanned by the last two dimensions. Stimulus-aligned neural geometry appeared in this subspace exclusively during the late delay period. Thus, the location cue transferred from the first two dimensions to the last two dimensions.

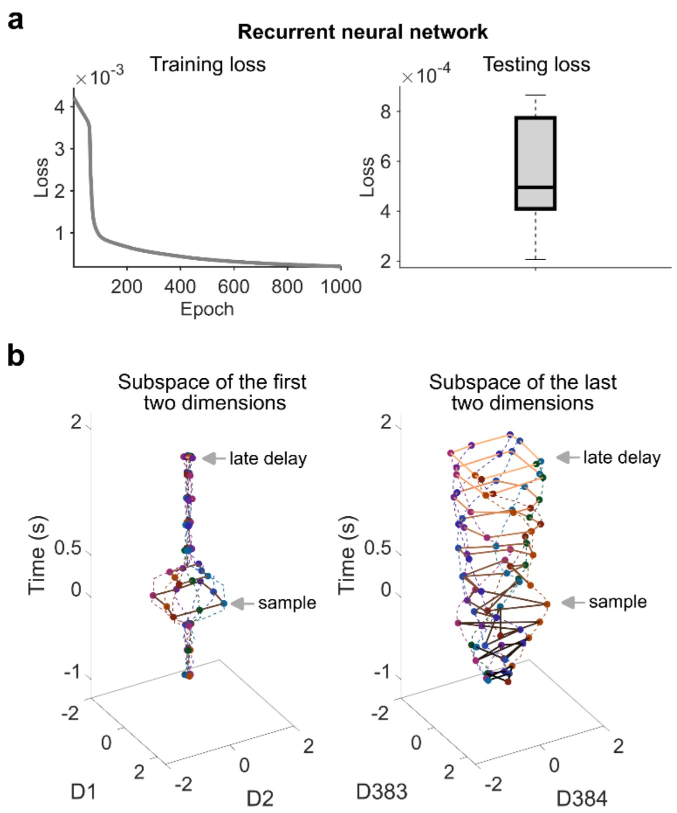


**Figure 9. A conventional RNN model executed the representation transfer task using a rotational mechanism.**

**(a)** Left: Over 1,000 training epochs, the loss decreased and converged to near zero. Right: When evaluated on additionally sampled test data, the test loss remained at a low level (median = 0.0005). **(b)** Left: The hidden states across the trial were projected onto the subspace spanned by the first two dimensions. As with the recurrent FWP model, stimulus-aligned neural geometry emerged transiently in this subspace during sample presentation but was absent following sample offset. Right: Stimulus-aligned neural geometry appeared in the subspace spanned by the last two dimensions after sample presentation and underwent continuous rotation within this subspace through the late delay period. This rotational dynamics differs from the transition dynamics observed in the empirical data.

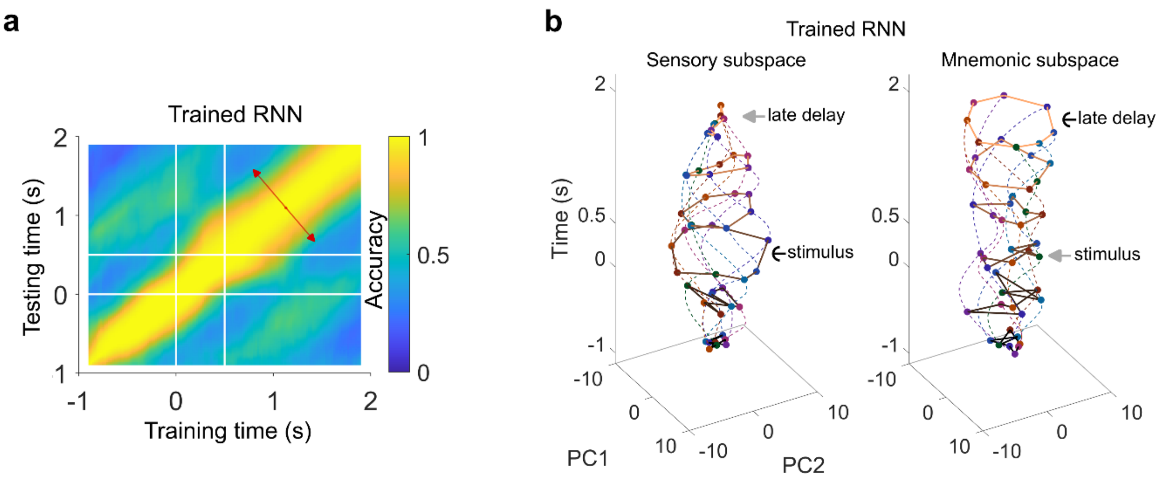


**Figure 10. The RNN model transformed its coding subspace through rotational dynamics.** This rotation was evident in both cross-temporal decoding and projection analyses. **(a)** The cross-temporal decoding accuracy matrix exhibited a grating-like pattern parallel to the diagonal. This pattern indicates that decoding accuracy fluctuated along the direction orthogonal to the diagonal (denoted by red arrows), suggesting periodic movement of the hidden states over time. **(b)** Rotation of the hidden states was clearly observed in both the sensory and mnemonic subspaces.

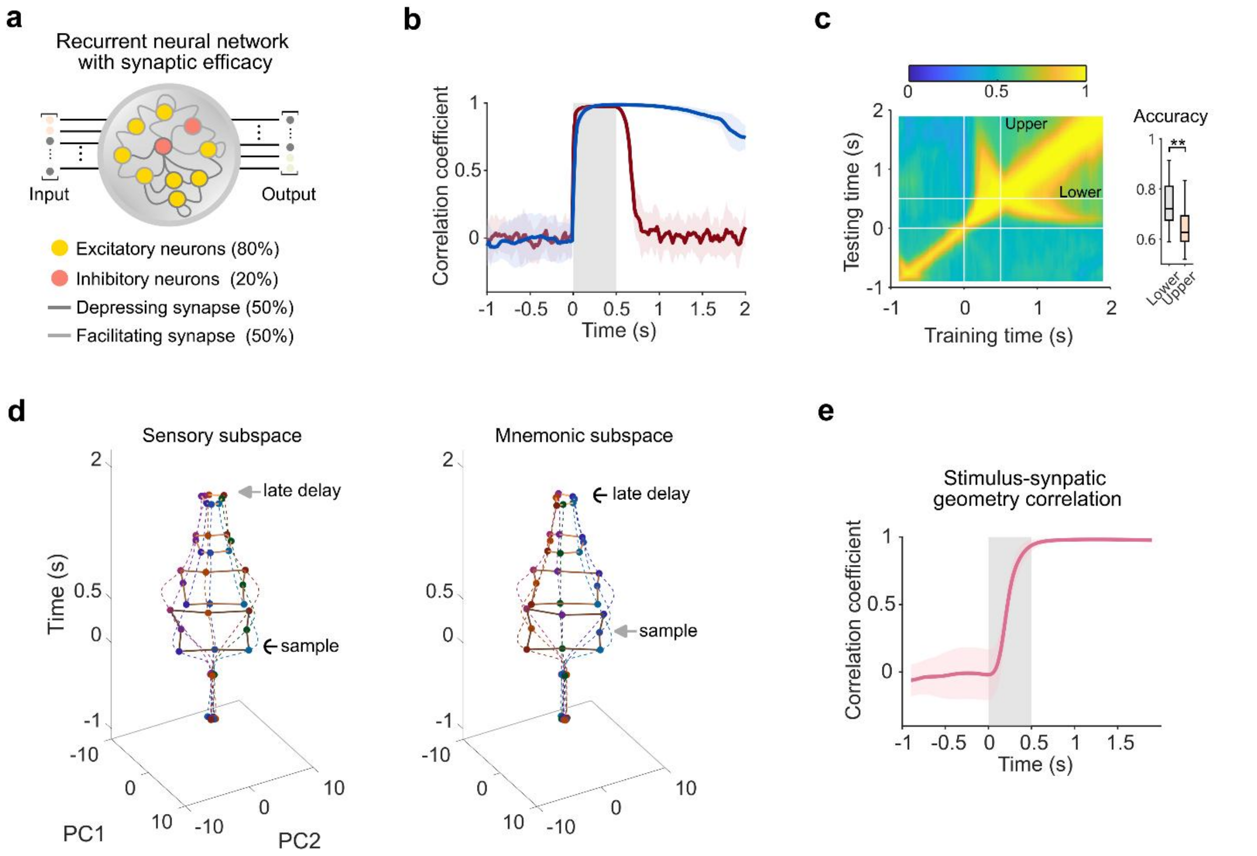


**Figure 11. A more biologically constrained RNN incorporating excitation-inhibition balance and synaptic-efficacy-gated recurrent connections.**

**a)** The extended RNN model in (Masse et al., 2019) added two features to vanilla RNNs: excitation/inhibition balance, with 80% excitatory and 20% inhibitory neurons; short-term synaptic plasticity (Mongillo et al., 2008), including half facilitating and half depressing synapses. **b)** Upon stimulus onset, the correlation rapidly rose to values close to 1. It then slowly decreased to a moderate level by the end of the delay. **c)** Left: The decoders trained in the stimulus period generalized well in the early-middle delay period. Right: the accuracies in the lower-right triangular region of the delay-period submatrix were significantly higher (t-test, $p<0.001$), opposite to empirical observation. **d)** In both the sensory and mnemonic subspaces, stimulus-aligned neural geometry appeared upon sample presentation and gradually reduced its size. **e)** Because the recurrent connections had plasticity, we also evaluated synaptic states of this model using the same method in (Masse et al., 2019). The stimulus-synaptic geometry rose to 1 during sample presentation and maintained at this value throughout the delay period, indicating that location cues were stored in the synaptic states. Gray region denotes sample presentation.

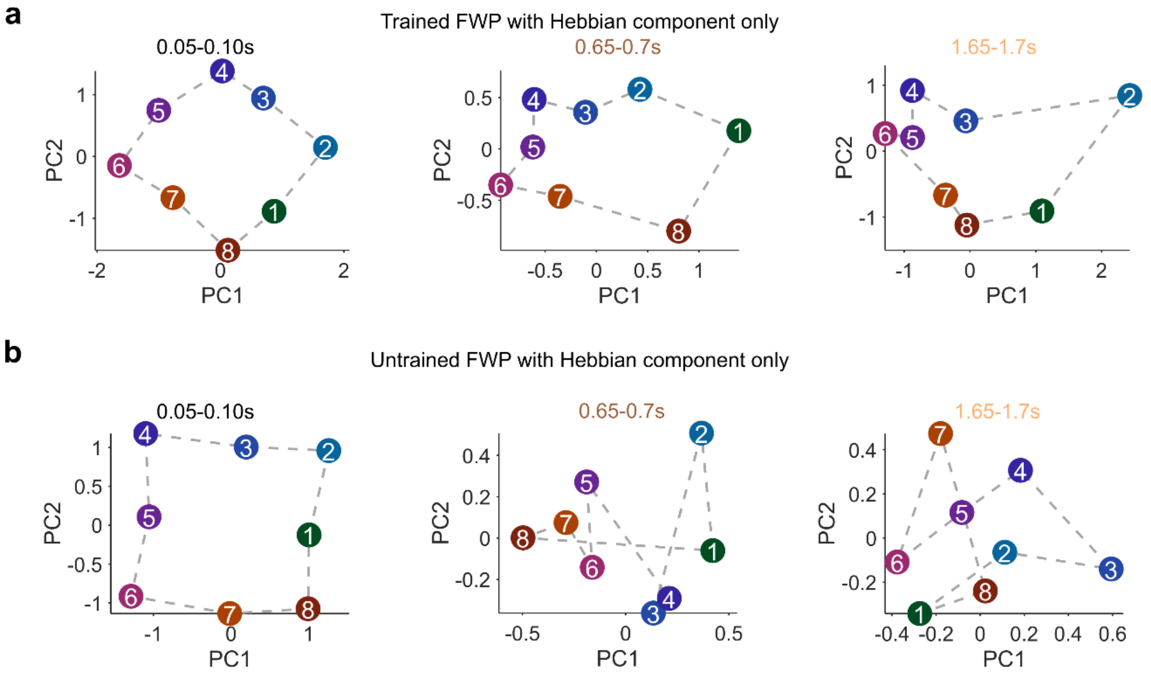


**Figure 12. For recurrent FWP models employing only the Hebbian component, stimulus information remained continuously readable from neural activity.**

Hidden states were projected onto the top two principal components at representative time points during the sample, early delay, and late delay periods. **(a)** In the trained model, neural geometries in the sample, early delay, and late delay periods all remained aligned with the stimulus geometry. **(b)** In the untrained model, neural geometries following sample presentation did not align with the stimulus geometry.

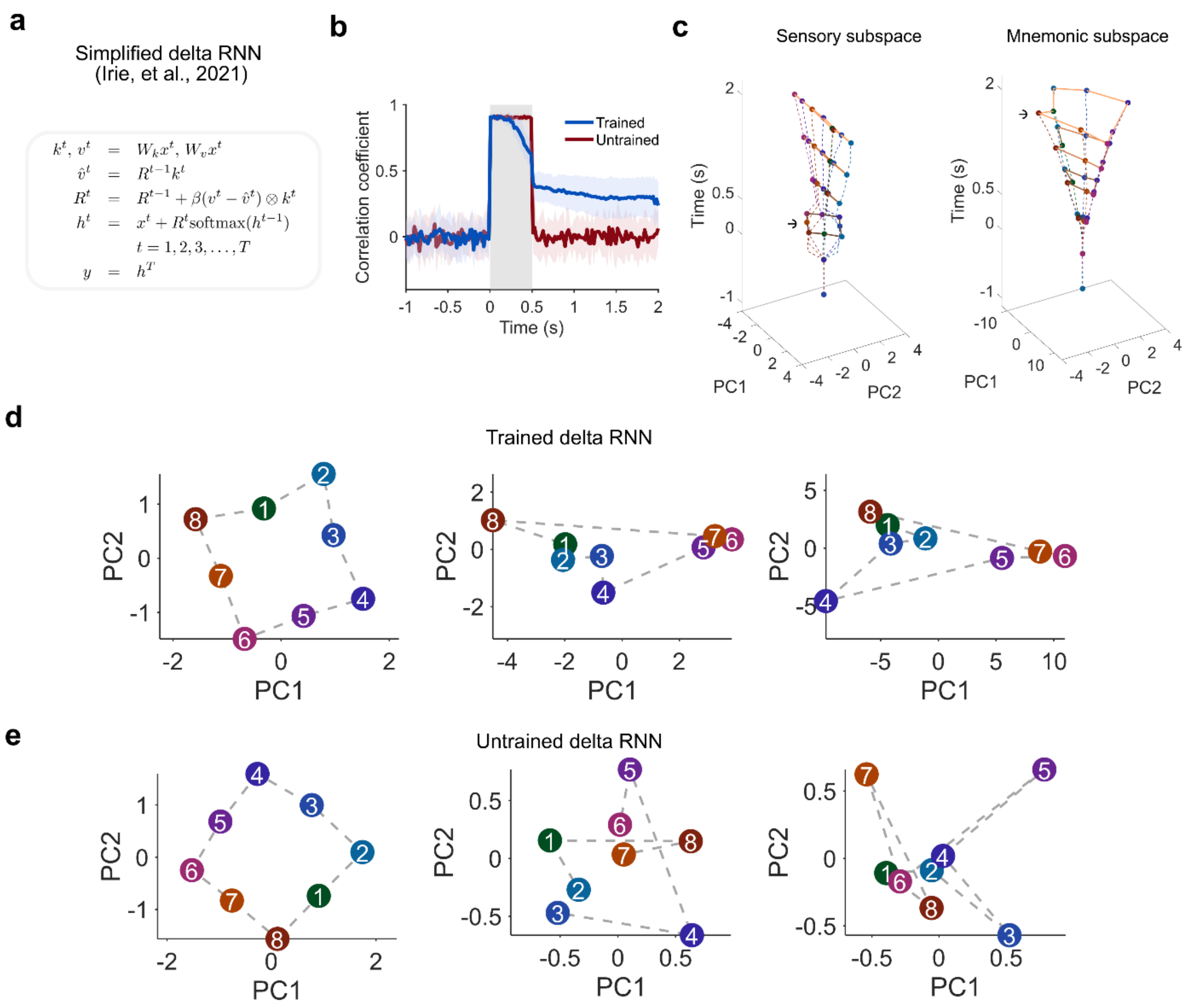


**Figure 13. A recurrent FWP model employing only the Hebbian component but using a delta-rule-style fast-weight update (Irie et al., 2021) relied on persistent activity to maintain memory.**

**(a)** Description of the delta-rule-style fast-weight update rule. **(b)** The time-resolved stimulus-neural geometry correlation of the trained model rose to 1 during sample presentation, declined to a moderate level following sample offset, and remained at this level throughout the delay period. This pattern resembles that of the recurrent FWP model using the simple Hebbian-only update rule. **(c)** Left: Stimulus-aligned neural geometry was transiently present during sample presentation. Right: Following stimulus offset, partial stimulus-aligned neural geometry emerged in the mnemonic subspace and expanded over time. **(d)** In the trained model, neural geometries in the sample, early delay, and late delay periods were all stimulus-aligned, although the fidelity of alignment decreased during the delay period. **(e)** In the untrained model, neural geometries following sample presentation did not align with the stimulus geometry.

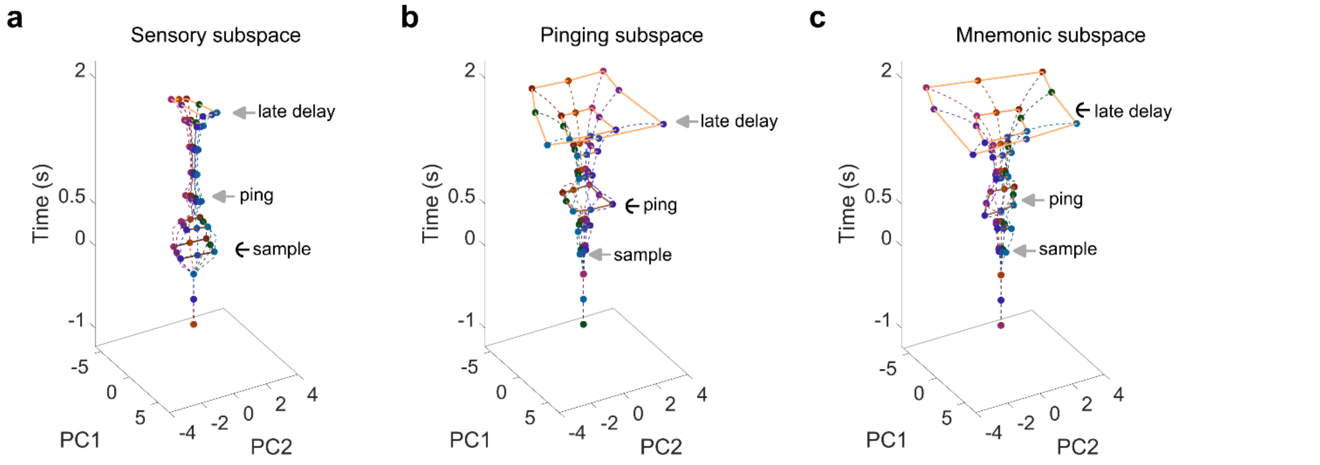


**Figure 14. Pinging-evoked representation was expressed in the mnemonic subspace.** **(a)** Hidden states during the pinging period exhibited minimal projection onto the sensory subspace. **(b)** Hidden states across the trial were projected onto the subspace spanned by the top two principal components of activity during the pinging period (pinging subspace). Hidden states in the sample period showed a small projection onto the pinging subspace, whereas late-delay states exhibited a large projection. **(c)** The projected neural geometries of both the pinging-period states and the late-delay states had magnitudes comparable to those in the pinging subspace, indicating that the pinging subspace and the mnemonic subspace largely overlapped.

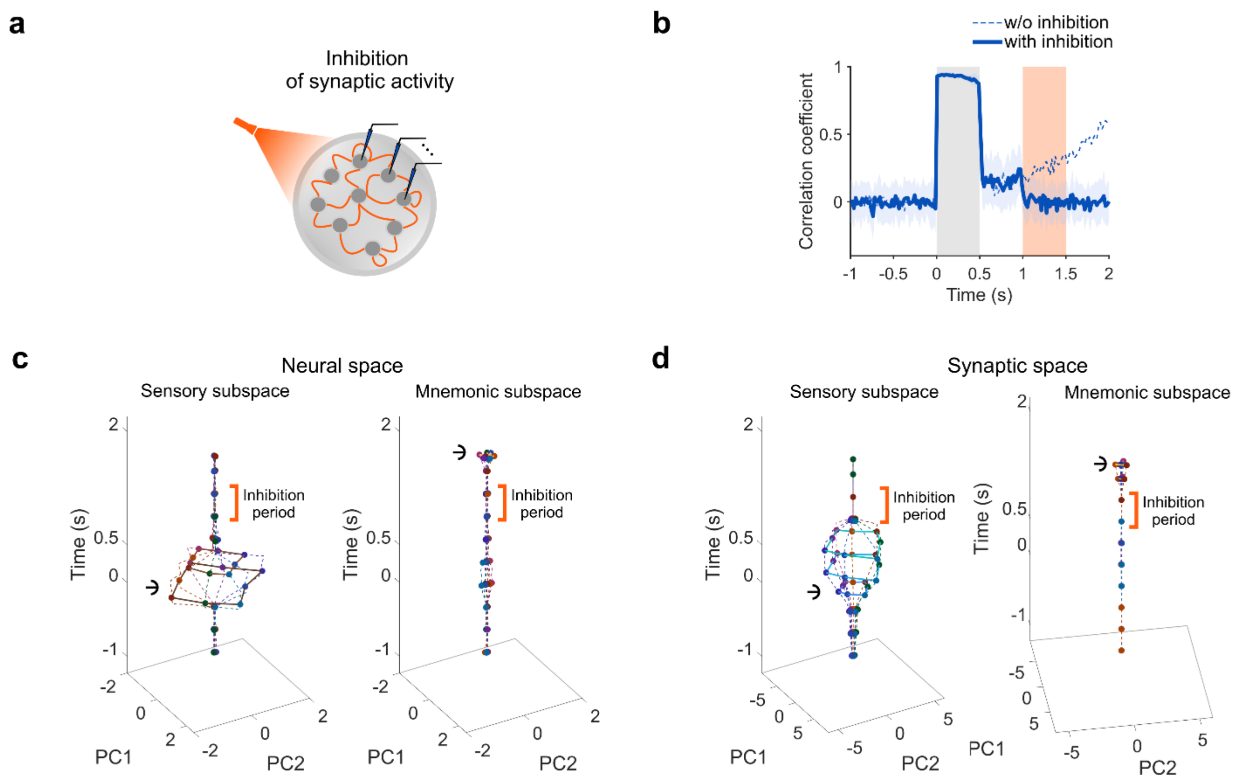


**Figure 15. When the stored information in the recurrent weights was eliminated, the ramping trajectory immediately collapsed to baseline.**

**(a)** We simulated optogenetic inhibition of synaptic activity by setting each element of the recurrent weight matrix to a value independently sampled from a zero-mean Gaussian distribution with a standard deviation of $1\times10^{-10}$, thereby erasing the stored memory. **(b)** The resulting time-resolved stimulus–neural geometry correlation dropped to zero immediately upon inhibition onset. Gray region denotes sample presentation. Orange region denotes the inhibition period. **(c)** Cross-temporal projection analysis revealed that no stimulus-aligned neural geometry emerged in the mnemonic subspace following inhibition. **(d)** Cross-temporal projection analysis of the synaptic states further demonstrated that the synaptic geometry collapsed upon inhibition.

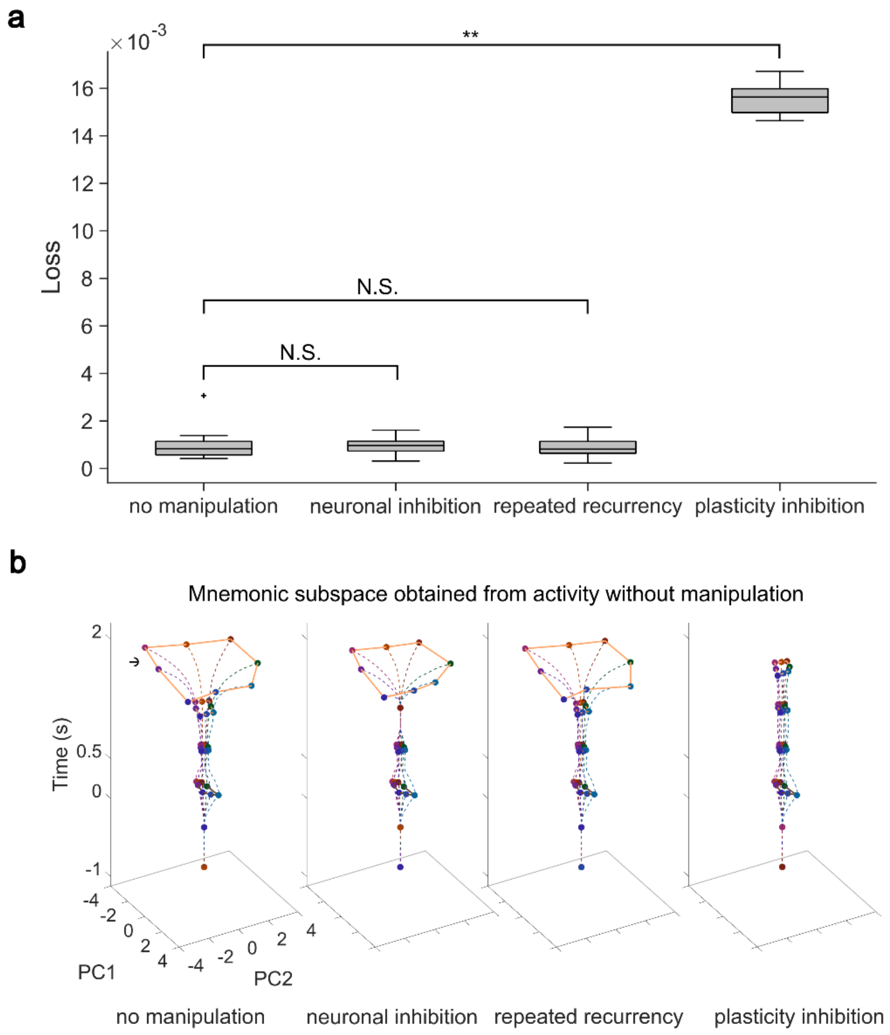


**Figure 16. The neural states transferred to the mnemonic subspace after inhibition of neuronal activity as though no perturbation was applied.**

**(a)** Testing loss of the recurrent FWP model under four conditions: no manipulation, neuronal activity inhibition, neuronal inhibition with repeated recurrent computation, and inhibition of synaptic plasticity. A one-way ANOVA revealed a statistically significant effect ($F(3,76) = 4407.6, p < 0.001$). Multiple comparisons with Bonferroni correction showed that the testing loss in the neuronal inhibition and repeated recurrency conditions did not differ significantly from the no-manipulation condition ($p = 1.0$), whereas the testing loss in the plasticity inhibition condition differed significantly from the no-manipulation condition ($p < 0.001$). **(b)** We defined the mnemonic subspace using the late-delay hidden state activity from the no-manipulation condition and projected the hidden states from the neuronal inhibition, repeated recurrency, and plasticity inhibition conditions onto this subspace across time. The mnemonic neural geometries in the no-manipulation, neuronal inhibition, and repeated recurrency conditions exhibited similar magnitudes, whereas the mnemonic neural geometry in the plasticity inhibition condition was substantially smaller. These results indicate that perturbation of neuronal activity had no effect on the mnemonic states, whereas plasticity inhibition did.

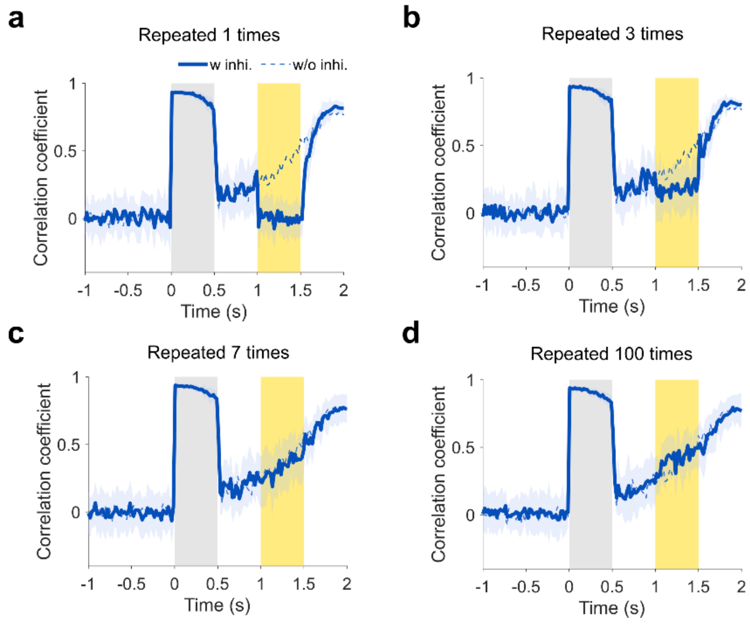


**Figure 17. Recovery of the ramping trajectory with increased repetition of recurrent computation.**

At each time step during the neuronal activity inhibition period, the hidden state activity of the fast net was initially clamped near zero. While the recurrent weight matrix remained fixed, the hidden state was allowed to pass through the recurrent weight matrix multiple times, enabling repetitive recurrent computation. **(a)** With a single repetition of recurrent computation, the trajectory of the stimulus-neural geometry correlation remained at baseline throughout the inhibition period. **(b)** With three repetitions, the correlation during the inhibition period rose to an elevated level but remained below the uninhibited trajectory. **(c)** With seven repetitions, the correlation trajectory overlapped with that observed in the absence of inhibition. **(d)** Beyond seven repetitions, the correlation trajectory did not increase further; instead, it remained close to the uninhibited trajectory.

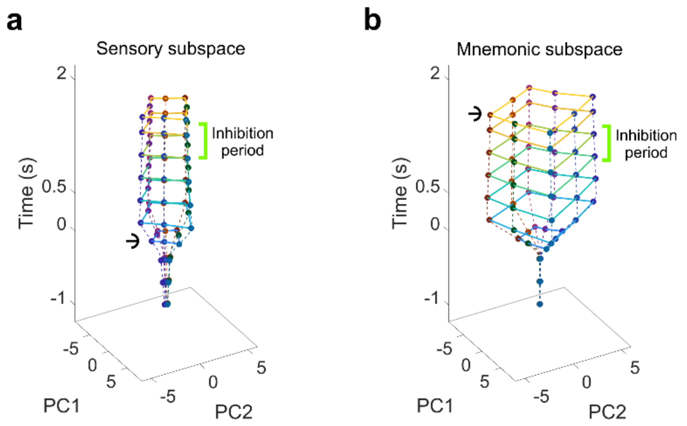


**Figure 18. The stimulus-synaptic geometry remained intact during the update-blockade period.**

**(a)** The sensory subspace was defined by the synaptic states during sample presentation. Stimulus-aligned synaptic geometry emerged immediately upon sample onset and remained stable thereafter. Blockade of fast-weight updates had no effect on the synaptic geometry. **(b)** Same as (a), except that the mnemonic subspace was defined by the synaptic states during the late delay period. Blockade of fast-weight updates likewise did not alter the stable synaptic geometry.

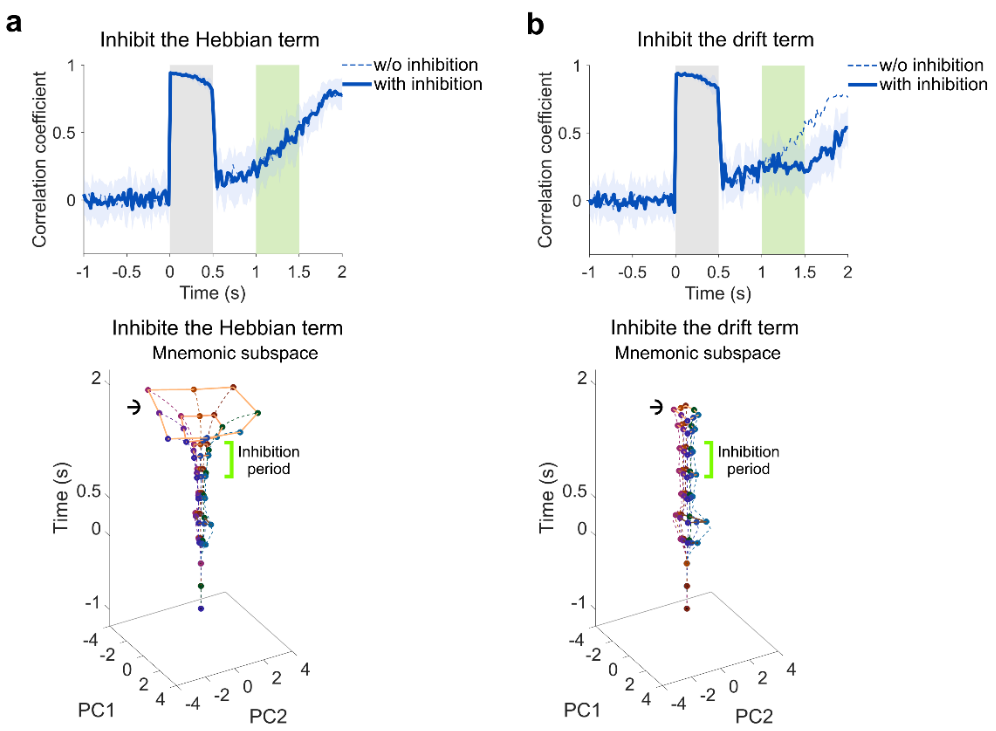


**Figure 19. The retrieval process was primarily driven by the drift component in the fast-weight update rule.** The fast-weight update rule consisted of a Hebbian component and a drift component. Having shown that blocking fast-weight updates disrupted the ramping trajectory of the stimulus-neural geometry correlation, we next investigated which component was the primary driver of this effect. **(a)** During the inhibition period, the Hebbian component of the fast-weight update rule was silenced. Top: The resulting trajectory of the stimulus-neural geometry correlation overlapped with that observed in the absence of inhibition. Bottom: The neural geometry in the mnemonic subspace was unaffected by this inhibition. The green region denotes the synaptic plasticity inhibition period. **(b)** During the inhibition period, the drift component was silenced. Top: The resulting trajectory deviated from the uninhibited trajectory. It ceased increasing during the inhibition period and, although it resumed increasing after inhibition was lifted, it reached only a lower value by the end of the delay. This pattern resembled that produced by blocking the entire update signal. Bottom: The neural geometry in the mnemonic subspace was markedly affected by the inhibition, exhibiting a substantially smaller magnitude than that observed without inhibition.

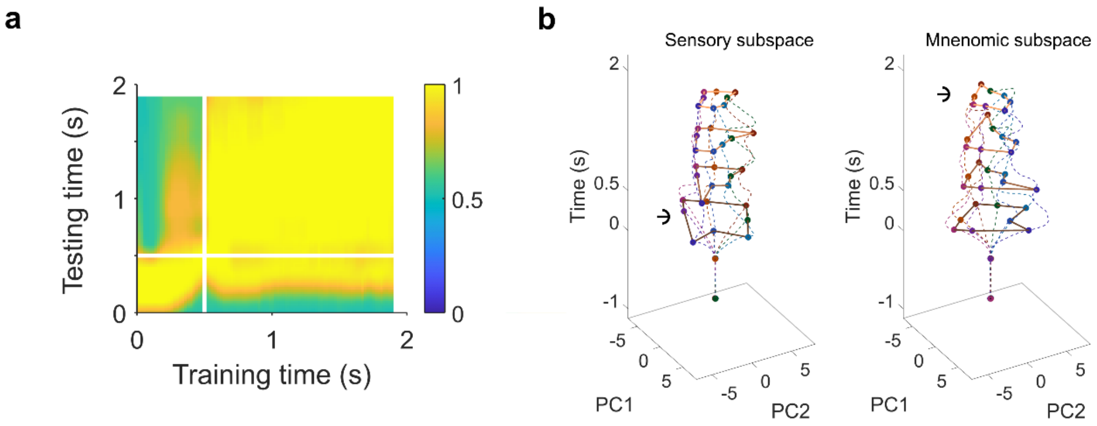


**Figure 20. Cross-temporal analysis of the recurrent FWP model trained on the variable-delay task revealed a more stable delay-period code.**

**(a)** Decoders trained at a single time point during the delay period generalized well to other delay-period time points. **(b)** In both the sensory and mnemonic subspaces, stimulus-aligned neural geometry emerged after sample onset, persisted throughout the delay period, and exhibited no rotational dynamics.

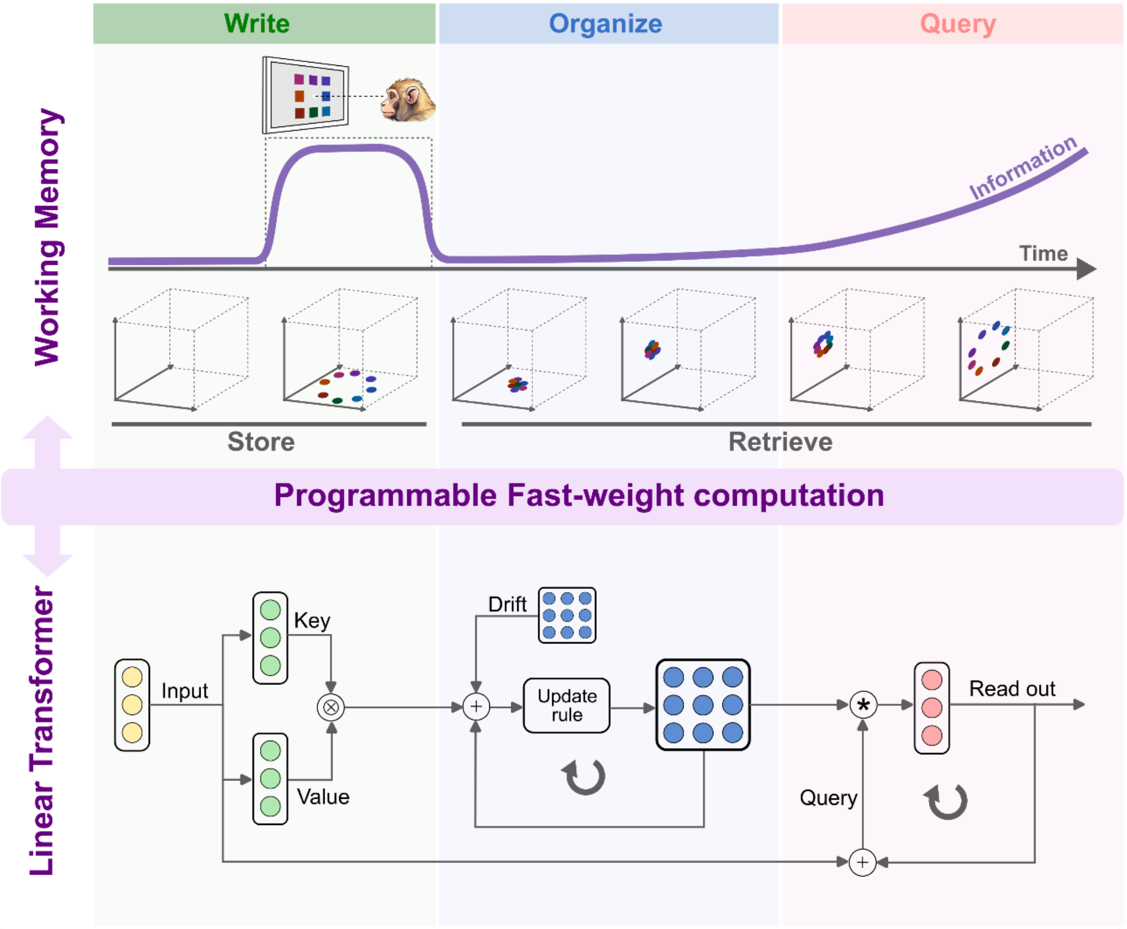


**Figure 21. Biological working memory can be interpreted through an algorithmic principle that is shared with linear Transformer.**

Top: In a single trial of the delayed task, stimulus information is encoded within one neural state subspace and stored as latent memory during sample presentation. During the subsequent delay period, this information is gradually retrieved into a distinct neural state subspace. The dashed box marks the sample presentation period. Bottom: This neural process corresponds to the computational steps of a linear Transformer architecture. Key and value vectors derived from the input implement memory writing. A writing signal and a drift signal recursively update the fast-weight matrix to organize the stored memory. A query signal, composed of the input and hidden state, iteratively reads out the memorized information in the desired format. These correspondences reflect that both working memory and linear transformer could be understood as programmable fast-weight computation.

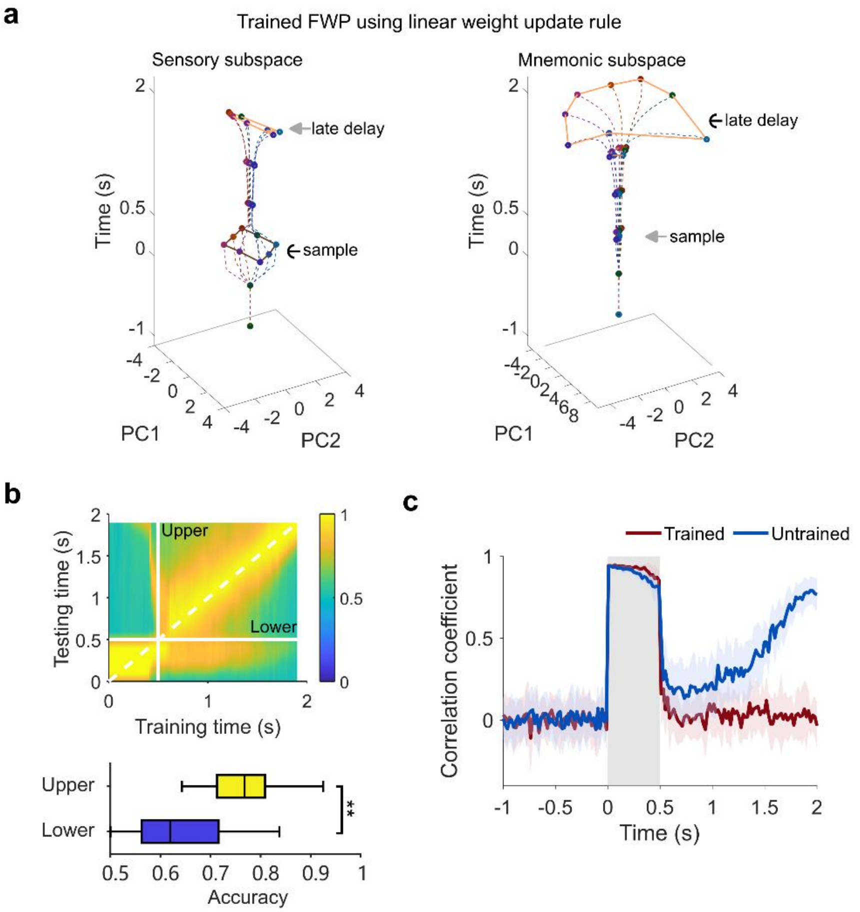


**Figure 22. A recurrent FWP model with linear weight updating behaved similarly to the model employing the hyperbolic tangent function.**

**(a)** Cross-temporal projections of fast-net activity into the sensory subspace (left) and the mnemonic subspace (right) after model training. Stimulus-period activity was prominently expressed in the sensory subspace, whereas late-delay activity was prominently expressed in the mnemonic subspace. **(b)** Cross-temporal decoding matrix for fast-net activity. White lines indicate stimulus offset; the dashed line indicates the matrix diagonal. "Upper" and "Lower" denote the upper-left and lower-right triangular regions of the delay-period submatrix, respectively. The bottom panel displays decoding accuracies in the upper and lower triangular regions; the upper triangular region exhibited higher accuracies. **(c)** Time-resolved stimulus-neural geometry correlation for trained and untrained models. The gray region denotes sample presentation, and colored shading indicates the standard deviation across 20 repetitions. **: $p < 0.001$.
